# Microglial ERK signaling is a critical regulator of pro-inflammatory immune responses in Alzheimer’s disease

**DOI:** 10.1101/798215

**Authors:** Michael J Chen, Supriya Ramesha, Laura D. Weinstock, Tianwen Gao, Linyang Ping, Hailian Xiao, Eric B Dammer, Duc D Duong, Allan I Levey, James J Lah, Nicholas T Seyfried, Levi B. Wood, Srikant Rangaraju

## Abstract

**Background:** The mitogen-activated protein kinase (MAPK) pathway is a central regulator of gene expression, pro-survival signaling, and inflammation. However, the importance of MAPK pathway signaling in regulating microglia-mediated neuroinflammation in Alzheimer’s Disease (AD) remains unclear. Here we examined the role of MAPK signaling in microglia using pre-clinical *in-vitro* and *in-vivo* models of AD pathology integrated with quantitative proteomics studies of post-mortem human brains.

**Methods:** We performed multiplexed immunoassay analyses of MAPK phosphoproteins, particularly ERK1/2, in acutely-isolated microglia and brain tissue from wild-type and 5xFAD mice. Neuropathological studies of mouse and human brain tissues were performed to quantify total and phosphorylated ERK protein in AD. The importance of ERK signaling in unstimulated and interferon γ (IFNγ)-stimulated primary microglia cultures was investigated using NanoString transcriptomic profiling, coupled with functional assays of amyloid β (Aβ) and neuronal phagocytosis. Receptor tyrosine kinases (RTKs) likely responsible for ERK signaling in homeostatic microglia and disease-associated-microglia (DAM) states and ERK-regulated human AD risk genes were identified using gene expression data. Total and phosphorylated MAPKs in human post-mortem brain tissues were measured in quantitative proteomic datasets.

**Results:** Phosphorylated ERK was the most strongly up-regulated signaling protein within the MAPK pathway in microglia acutely isolated from 5xFAD brains. Neuroinflammatory transcriptomic and phagocytic profiling of mouse microglia confirmed that ERK is a critical regulator of IFNγ-mediated pro-inflammatory activation of microglia, although it was also important for constitutive microglial functions. Phospho-ERK was an upstream regulator of disease-associated microglia (DAM) gene expression (Trem2, Tyrobp), as well as of several human AD risk genes (Bin1, Cd33, Trem2, Cnn2). Among RTKs that signal via ERK, CSF1R and MERTK were primarily expressed by homeostatic microglia while AXL and FLT1 were likely regulators of ERK signaling in DAM. Within DAM, FLT4 and IGF1R were specifically expressed by pro- and anti-inflammatory DAM sub-profiles respectively. In quantitative proteomic analyses of post-mortem human brains from non-disease, asymptomatic and cognitively-impaired AD cases, ERK1 and ERK2 were the only MAPK pathway signaling proteins with increased protein expression and positive associations with neuropathological grade. Moreover, in a phospho-proteomic study of post-mortem human brains from controls, asymptomatic and symptomatic AD cases, we found evidence for a progressive increased flux through the ERK signaling pathway.

**Conclusions:** Our integrated analyses using pre-clinical models and human proteomic data strongly suggest that ERK phosphorylation in microglia is a critical regulator of pro-inflammatory immune response in AD pathogenesis and that modulation of ERK via upstream RTKs may reveal novel avenues for immunomodulation.

## INTRODUCTION

Neuroinflammation is now well recognized to play a critical role in the pathogenesis of Alzheimer’s disease (AD). As the primary immune cells of the brain, microglia are the key enactors of neuroinflammation in AD (Hickman et al, 2018; Lambert et al, 2013). However, microglia can mediate beneficial and detrimental neuroinflammatory mechanisms during disease. Indeed, because microglia are transcriptionally diverse and highly responsive to their environment (Dubbelaar et al, 2018), they contextually respond to acute and chronic tissues disease states determined by both the primary disease mechanism and disease duration (Bachiller et al, 2018; Tay et al, 2017). In general, detrimental consequences of microglial responses in AD are mediated by pro-inflammatory mechanisms, which include release of neurotoxic molecules (ROS, NO), production of cytokines (IL1β, TNFα, IL6), phagocytosis of live neurons or of healthy synapses and dendrites, impaired phagocytic clearance of debris and pathological protein aggregates. Microglial activity can also promote pathogenesis via indirect mechanisms, such as dysregulation of astrocytic or oligodendroglial functions, which then impact neuronal health and survival (Block et al, 2007; Geraghty et al, 2019; Hagemeyer et al, 2017; Liddelow et al, 2017; Sellgren et al, 2019). Selective inhibition of these pro-inflammatory responses without impacting homeostatic functions is the desired immune consequence of neuro-immunomodulatory therapies for AD. Therefore, defining critical regulators of homeostatic, disease-associated and pro-inflammatory microglial responses in AD is necessary for the development of effective neuro-immunomodulatory therapies for AD (Rangaraju et al, 2018b).

Distinct microglial states and their immunological profiles are likely to be determined by upstream signaling events that are engaged when microglia are exposed to protein signals, cytokines, pathological proteins and oxidative stress in the setting of neuronal injury or neurodegeneration. Signaling pathways, such as the mitogen activated protein kinase (MAPK) family of proteins, are engaged very early in immune cells following signals that activate receptor tyrosine kinases (RTKs) (Cargnello & Roux, 2011). The MAPK pathway includes over 60 proteins which can be broadly categorized into three families, namely the ERK, JNK, and p38 MAPK cascades (Yates et al, 2017). These dynamic cascades are initiated via activation of a cell surface receptor, which triggers serial phosphorylation events by serine/threonine protein kinases, ultimately activating or suppressing transcriptional factors which in turn regulate gene expression (Arkun & Yasemi, 2018; Santos et al, 2007). ERK signaling is initiated by RTKs that are typically receptors for growth factors. JNK signaling is activated by cellular stress and p38 is activated by inflammatory stimuli and cytokines. There is also significant cross talk between these three cascades, particularly JNK and p38 (Zhang & Liu, 2002). While these signaling events are typically transient and often necessary for homeostasis, constitutive functions, and proliferation, sustained activation may lead to dysfunction and chronic inflammatory activation in microglia (Kim et al, 2004). However, the importance of MAPK signaling in regulating microglia-mediated neuroinflammation in AD remains unclear.

In this manuscript, we investigate the role of MAPK signaling in microglia using multiplexed immunoassay phospho-protein signaling studies of concomitant analysis of acutely-isolated microglia and whole-brain tissues from 5xFAD mice. Our data for the first time reveal a microglial-specific activation of ERK signaling in a pre-clinical model of AD pathology. We also provide evidence that ERK proteins are highly abundant and functionally important MAPKs in microglia. In neuroinflammatory transcriptomic profiling of primary mouse microglia, we show that ERK signaling is a critical regulator of pro-inflammatory microglial activation in response to interferon γ (IFNγ), in addition to being an upstream regulator of several disease-associated microglial genes as well as genetic risk factors for late-onset AD in humans. We further interrogated human post-mortem quantitative proteomics data from control and AD brains and provide evidence for increased activation of the ERK cascade in human AD. Using this integrated analysis of microglial cultures, an AD mouse model, and human AD tissues, we provide novel insights into the role of ERK signaling in microglia as a critical regulator of pro-inflammatory immune responses and causal immune mechanisms of AD pathogenesis.

## MATERIALS AND METHODS

### Animals

C57BL/6J and 5xFAD mice used for the studies were housed in the Department of Animal Resources at Emory University under standard conditions with no special food/water accommodations. Institutional Animal Care and Use Committee approval was obtained prior to *in-vivo* work and all work was performed in strict accordance with the Guide for the Care and Use of Laboratory Animals of the National Institutes of Health.

### Reagents

The ERK1/2 inhibitor SCH772984 was obtained from Selleckchem (S7101) (Morris et al, 2013). The following antibodies were used for flow cytometry: CD45-PE-Cy7 Rat mAb (BD Biosciences, #552848), LIVE/DEAD Fixable Blue Dead Cell Stain (Invitrogen, L34961), CD11b-APC-Cy7 Rat mAb (BD Biosciences, #557657), phospho-ERK1/2 Rabbit pAb (Cell Signaling Technology, #9101S), ERK1/2 Rabbit pAb (R&D Systems, AF1576) and for immunohistochemistry: ERK1/2 Rabbit pAb (R&D Systems, AF1576), phospho-ERK1/2 Rabbit pAb (R&D Systems, AF1018).

### Mouse microglia isolation and in-vitro studies

Adult mouse microglia were isolated from the brains of wild-type and age-matched 5xFAD mice (age 6-7 mo, n=7/group, 3 males and 4 females per group) using protocols we have published previously (Rangaraju et al, 2018b). Briefly, mononuclear brain phagocytes were isolated by Percoll density centrifugation (35%/70% isotonic Percoll gradient) and then further enriched for CD11b+ microglia by MACS columns (Miltenyl Biotec). These were immediately washed, and pelleted cells were immediately frozen on dry ice and then lysed for Luminex phospho-protein signaling studies.

Primary microglia cultures were prepared from P0-3 post-natal mouse brains as previously described (Gao et al, 2019). Briefly, brain tissue was harvested from C57BL/6J mice (P0 to P3) and digested with Trypsin for 15 min. Trypsinization was halted by the addition Dulbecco’s Modified Eagle Medium: Nutrient Mixture F-12 (DMEM-F12) supplemented with 10% fetal bovine serum (FBS) and 1% penicillin-streptomycin-glutamine. Once filtered using a 40 µm nylon mesh strainer, the resulting cell suspension was positively selected for CD11b^+^ cells using mini-MACS columns. The cells, almost entirely consisting of microglia, were seeded into poly-L-lysine-coated wells and cultured in a humidified CO_2_ incubator at 37□. Following 24 h of incubation, half of the medium was replaced with fresh medium and 1µM SCH772984, a selective small molecule inhibitor of ERK 1/2 (IC_50_ for ERK1: 4nM, IC_50_ for ERK2: 1 nM) was added to the culture (Morris et al, 2013). After 1 h of pre-incubation, cells were stimulated with 1 ng/mL IFNγ (Sigma) and then incubated for 24 h, after which cells were trypsinized and harvested for downstream analyses.

### Luminex phospho-protein analyses of acutely-isolated microglia from mouse brain

Collected microglia were lysed in Bio-Plex lysis buffer per kit instructions (Bio-Rad) with the addition of 1 Complete Mini/5mL of buffer. Cells were lysed for >10 min on an end-over-end rotator, then centrifuged at 13kRPM for 10 min with supernatant collected and stored at −80^ο^C. Samples were normalized to total protein using Pierce bovine serum albumin (BSA) Protein Assay kit (ThermoFisher Scientific, Waltham, MA, USA) and normalized to 0.1µg in Bio-Plex lysis buffer. Phospho-signaling was quantified using the MAPK/SAPK Signaling 10-Plex Multiplex assay (Millipore Sigma, 48-660MAG). Phosphorylation sites for the MAPK/SAPK as follows: ATF2 (Thr71), ERK (Thr185/Tyr187), HSP27* (Ser78), JNK (Thr183/Tyr185), c-Jun (Ser73), MEK1 (Ser222), MSK1 (Ser212), p38 (Thr180/Tyr182), p53* (Ser15) and STAT1 (Tyr707). Analytes marked with an asterisk have not been reported to be mouse-reactive and were removed from the analysis. The assay was read out using a MAGPIX instrument (Luminex, Austin, TX, USA).

### NanoString transcriptomic profiling of microglia and statistical analyses

NanoString transcriptomic profiling of mRNA isolated from primary mouse microglia was performed as previously published (Gao et al, 2019). Primary microglia were treated with the ERK inhibitor in the presence or absence of IFNγ. The cells were washed in 1x Diethyl pyrocarbonate-phosphate buffered saline (DEPC-PBS) and lysed in TRIzol (Invitrogen). RNA was extracted from the lysates using the nCounter Low RNA Input Kit, and a quality control assessment was performed to determine concentration and integrity of RNA (RIN scores >8 for all samples). 100 ng of RNA per sample was run using the nCounter Mouse Neuroinflammation Panel (770 genes). The resulting gene expression data was imported into nSolver Analysis Software (version 4.0). A quality control check was done to assess the technical performance of the assay, and background thresholding parameters were established as the median of negative control counts. Only genes for which >50% of samples exceeded this threshold (n=678) were included in the final analysis. The expression of these genes was first normalized to the geometric mean of the positive controls and then normalized to the geometric mean of 12 housekeeping genes.

One-way analysis of variance (ANOVA) and Tukey’s Honestly Significant Difference (Tukey’s HSD) tests were performed for comparison across four groups for each transcript (R, version 3.5.1). Genes demonstrating differential expression across groups (ANOVA p<0.05) were included in K-means clustering analysis conducted using Morpheus (https://software.broadinstitute.org/morpheus) with a pre-defined cluster number of 5. Gene ontology (GO) enrichment analysis was then performed on the resulting clusters using GO-Elite (version 1.2.5), a software that identifies GO terms, KEGG pathway terms and Wikipathway terms that are enriched in lists of genes as compared to a reference list of genes (Zambon et al, 2012). In addition to K-means clustering, we also used t-weighted stochastic neighbor embedding (t-SNE) to visualize clusters of genes in two dimensions and determine whether known homeostatic microglia and disease-associated-microglia (DAM) genes mapped to specific gene clusters identified in our dataset. We used existing reference transcriptomes and network analyses to obtain lists of homeostatic and DAM genes (Keren-Shaul et al, 2017) and lists of genes specific to modules of co-expressed microglial genes that were identified using network analyses approaches (Rangaraju et al, 2018b).

### Immunohistochemistry (IHC) studies of ERK and phospho-ERK in mouse brain

We used fixed floating sections (20µm thick) of brain derived from wild-type/Cx3cr1-Yfp and 5xFAD/Cx3cr1-Yfp mice for immunofluorescence studies using established protocols described previously (Rangaraju et al, 2018a). The sections were blocked with 8% normal horse serum for 1 h and incubated with primary antibodies (anti-Phospho-ERK1/2, 1:20, 10ug/mL or anti-ERK1/2, 1:20, 10ug/mL, and anti-GFP 1:100) at 4 L overnight. Then, the sections were incubated with fluorophore-conjugated secondary antibodies for 1 h at room temperature (RT). Following 4 washes, the sections were mounted with hard mounting medium (VectorLabs, H1500) and DAPI for nuclear labeling. Negative controls (unstained sections and secondary alone sections) were performed for each experiment. Immunofluorescence imaging was performed using 20x and 60x (oil immersion) objectives on a fluorescence microscope (Olympus BX51 and Camera: Olympus DP70) using FITC, PE and DAPI filters. Image processing was performed using ImageJ software.

### Flow cytometry studies of microglia

Flow cytometric assays of microglia viability, amyloid β (Aβ) phagocytosis, and microglia-mediated neuronal phagocytosis were performed using *in-vitro* primary mouse microglia. To assess the viability of microglia under the experimental conditions used in the NanoString studies above, primary microglia were treated with the ERK inhibitor and/or IFNγ using the concentrations and durations described above. After 24 h of incubation at 37L, the cells were washed in 1x PBS and trypsinized for 15 min at 37□. The trypsin was quenched with DMEM with 10% FBS, and the cells were washed again in PBS and labelled with LIVE/DEAD Fixable Blue Stain in the dark at RT for 30 min. After washing with 2% normal horse serum and again with PBS, PE-Cy7 fluorophore-conjugated anti-mouse CD45 antibody was added to the microglia and allowed to incubate at RT in the dark for 30 min, after which the cells were washed in PBS. Compensation, gating, and data analysis for flow cytometry were performed according to previously established protocols (Rangaraju et al, 2018b). Viability was measured as the proportion of cells that were weakly fluorescent with the fixable blue dye.

The effect of ERK inhibition and inflammatory stimulation on microglial phagocytosis of fluorescent fibrillar amyloid-β_42_ (fAβ42) was determined through flow cytometry as previously described (Gao et al, 2019). Briefly, fAβ42 peptide conjugated to HiLyte Fluor 488 was mixed with 1% NH_4_OH and diluted in 1xPBS to achieve a concentration of 100µM, after which it was incubated in the dark for 6 days at RT. Once primary microglia had been treated with the ERK inhibitor and/or IFNγ, they were incubated with 2uM of fAβ42-488 for 1 h at 37□. The cells were collected and subsequently labelled with anti-CD45, after which flow cytometry was performed. Phagocytosis was assessed by observing the second peak of fluorescence emitted by phagocytic microglia, which was previously confirmed to represent actin-dependent phagocytosis (Rangaraju et al, 2018b).

To measure microglia-mediated neuronal phagocytosis, we performed microglia-neuron co-culture studies using mouse N2a neuroblastoma cells that were stably transduced to express GFP (Lenti-Ubc-GFP/FUGW backbone, Addgene #14883) and primary p0-3 mouse microglia and used flow cytometry to detect microglial uptake of GFP+ neuronal material as an indicator of phagocytosis of neurons. All GFP-N2a cells were washed twice with medium and allowed to grow to 70-80% confluence. 50,000 N2a cells were seeded into 12-well plates, and 24 h later, the medium was replaced with DMEM with 1% FBS and 10 µM retinoic acid to induce differentiation for 4 days (Namsi et al, 2018). Meanwhile, primary microglia were cultured in the presence or absence of IFNγ (1µg/mL) and treated with the ERK inhibitor (or DMSO control) at the same concentrations used in our transcriptomic studies. After 4 days of N2a neuronal differentiation, fresh medium with 1% FBS replaced the differentiation medium in which the N2a cells grew, and 100,000 microglia were seeded into each of the wells containing differentiated N2a cells. The co-culture was incubated overnight and then trypsinized for 2 min at 37□ to harvest the cells for antibody staining. The cells were washed in 1% FBS media and again in PBS and labelled with LIVE/DEAD indicator (fixable blue) and anti-CD45 (PC-Cy7). Microglial phagocytosis of neuronal cells was assessed by determining GFP positivity in microglia via flow cytometric analysis and appropriate negative controls (microglia without neurons). We compared the proportion of live CD45+ microglia that were also positive for GFP, across all the treatment conditions. In parallel, we also measured the proportion of live N2a-GFP+ cells in the co-culture by flow cytometry. These experiments were performed in biological triplicate.

Flow cytometric measurement of phospho-ERK1/2 expression was performed in acutely-isolated CNS mononuclear cells derived from wild-type and 5xFAD mice. Acutely-isolated CNS mononuclear cells were labeled with fluorophore-conjugated mAbs against CD11b (APC-Cy7) and CD45 (PE-Cy7), then fixed with fixation buffer for 10 min at RT. They were then permeabilized with cold methanol for 30 min at 4°C in the dark and blocked with Fc block (Biolegend Cat #553142) for 30 min. The cells were then incubated with primary anti-pERK1/2 (1:100) or anti-ERK (1:20) for 30 min followed by a secondary antibody (Goat anti-rabbit-FITC, 1:500) for 30 min and then taken for flow cytometric analyses. Appropriate negative (unstained), secondary-alone and fluorescence-minus-one (FMO) controls were performed for all experiments involving unconjugated primary antibodies. Live gated (based on FSC/SSC parameters) single cells (based on FSC-A and FSC-H parameters) that expressed CD11b and intermediate CD45 were included in the analysis and pERK1/2 expression was compared between wild-type and 5xFAD microglia.

### Quantitative reverse transcriptase PCR (qRT-PCR)

qRT-PCR validation studies were performed using RNA isolated from primary microglia that were cultured in the presence or absence of ERK inhibitor and IFNγ to mirror the conditions used in NanoString studies, using protocols described previously (Gao et al, 2019). RNA was isolated from microglia, and RNA concentration and purity were determined using the NanoDrop 2000 spectrophotometer. cDNA was synthesized, and qRT-PCR was performed on the 7500 Fast Real-time PCR System (Applied Biosystems) using cDNA, TaqMan PCR master mix (Applied Biosystems), and gene-specific TaqMan probes (Applied Biosystems) against Apoe (Mm01307193_g1), Tnf (Mm01210732_g1), Trem2 (Mm04209424_g1), Tyrobp (Mm00449152_m1), Spp1 (Mm00436767_m1), and Gapdh (Mm99999915_g1). All primer sets were run in duplicate for every RNA sample and 4 biological replicates were performed per condition. Gene expression was normalized to the housekeeping gene, GAPDH, and relative gene expression was calculated using the 2-ΔΔCt method.

### Human post-mortem brain quantitative proteomics data

Two post-mortem brain proteomic datasets were used for analyses, with cases obtained from the Goizueta Alzheimer’s Disease Research Center at Emory. The first is a published dataset in which dorsolateral prefrontal cortex from 10 non-disease controls, 10 AD cases, 10 with Parkinson’s disease (PD) and 10 with both AD and PD pathological features were sampled for tandem mass tag (TMT) proteomics (Ping et al, 2018) and the patient and pathological features as well as expression data are publicly available at http://proteomecentral.proteomexchange.org/cgi/GetDataset?ID=PXD007160.

The second dataset was a phospho-proteomics expression dataset of 6 non-disease controls, 6 asymptomatic AD cases and 6 AD cases with dementia. Methodology used for phospho-peptide enrichment by immobilized metal-affinity chromatography (IMAC) as well as isobaric tandem mass tag mass spectrometry have been previously published (Dammer et al, 2015; Johnson et al, 2018). The patient characteristics, pathological features, TMT mass spectrometry methodology and raw expression data pertaining to these samples are made available at https://www.synapse.org/#!Synapse:syn20820053/wiki/596078 and http://proteomecentral.proteomeexchange.org. For this manuscript, only phospho-peptide data pertaining to any of the MAPK family members were used for analyses and have been included in the supplemental data in this manuscript. All other data and analyses will be published separately.

### Other statistical considerations

GraphPad Prism (Ver. 5) and SPSS (Ver. 26) were used to create graphs and for statistical analyses. Error bars in all bar graphs represent the standard error of the mean. For experiments with >□2 groups, one-way ANOVA was performed to detect differences across groups and post-hoc pairwise comparisons were performed using Tukey’s HSD test. Statistical significance was set at *p* value <0.05 for all experiments.

## RESULTS

### Phosphorylated ERK and p38 MAPK is increased in microglia isolated from 5xFAD mice

MAPKs are comprised of three broad sub-families of signaling proteins, namely ERK, JNK and p38 pathways (Figure 1a). We hypothesized that chronic microglial activation in models of neurodegeneration may be characterized by signaling via distinct MAPK pathways. To determine whether MAPK signaling is chronically up-regulated in microglia in a mouse model of AD pathology, we used magnetic associated cell sorting (MACS) columns to acutely isolate CD11b+ brain myeloid cells from adult (6-7 mo old) wild-type and age- and sex-matched 5xFAD mice that exhibit accelerated accumulation of Aβ pathology in the brain. CD11b^+^ brain myeloid cells are comprised primarily (>95%) of microglia and a small proportion of CNS-infiltrating and perivascular blood-derived monocytes/macrophages (Rangaraju et al, 2018b). To broadly interrogate phospho-signaling within the MAPK pathway in CD11b+ microglia, we lysed the isolated microglia and used a Luminex multiplexed immunoassay (Millipore) quantify phosphorylation of 9 proteins within the MAPK pathway (Figure 1b). We simultaneously quantified signaling within whole brain samples obtained from the frontal cortex (Figure 1b) where Aβ pathology is abundant in 5xFAD mice. As compared to microglia isolated from WT mice, 5xFAD mouse microglia exhibited increased activation of ERK signaling, indicated by increased pERK1/2 and trend towards increased levels of pMEK1, an upstream ERK activator. Increased p38 MAPK activation was also observed indicated by higher p-p38 levels in 5xFAD microglia (Figure 1c). Unlike microglia, we did not observe any differences in MAPK signaling in whole brain (Figure 1d), suggesting that there are specific differences in microglial signaling in 5xFAD mice that are not broadly exhibited across the brain. These findings provide evidence for increased activation of specific MAPKs in microglia in 5xFAD mice.

**Figure 1.**
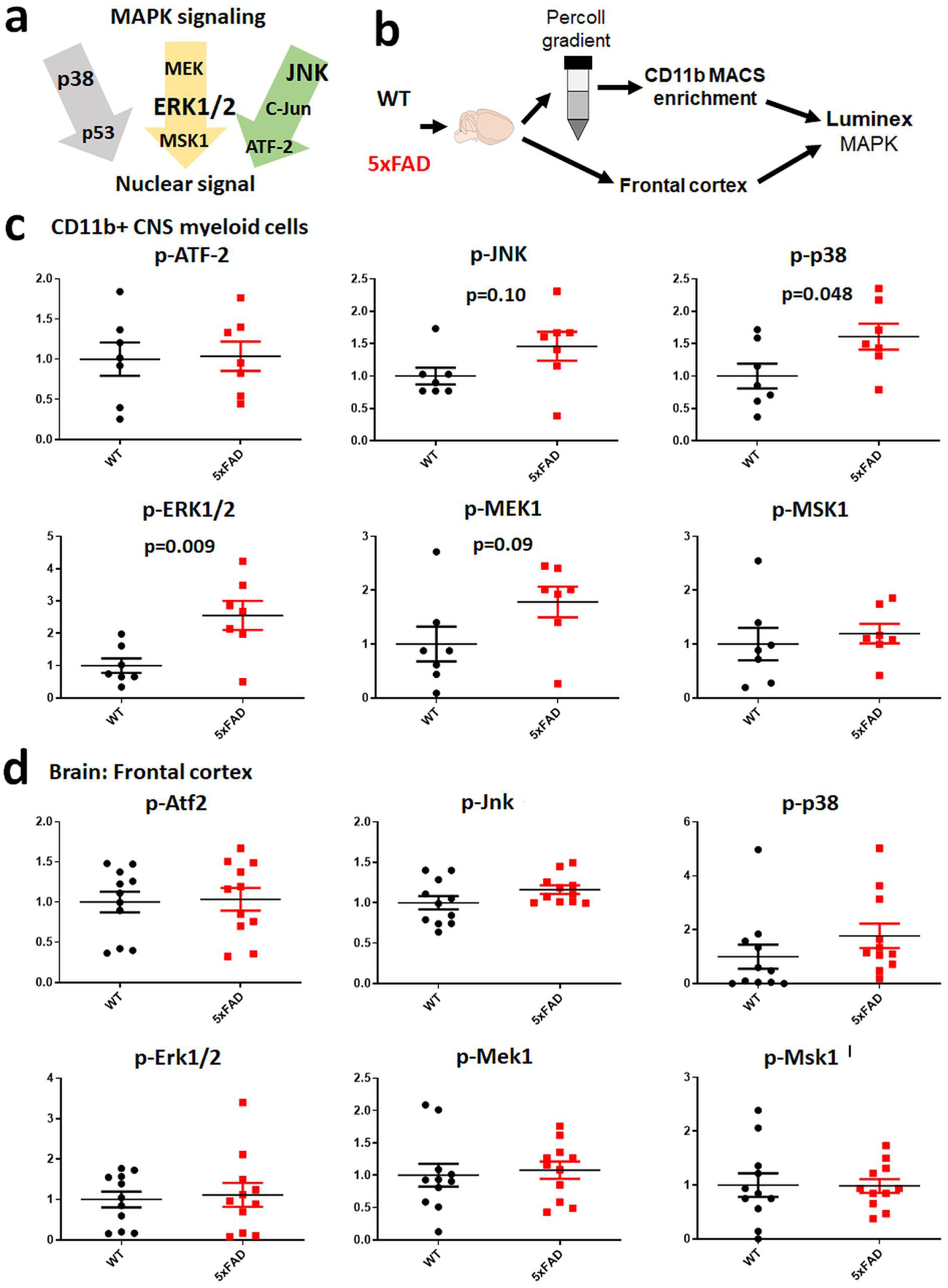
Increased activation of ERK1/2 and p38 MAPK signaling in microglia in mouse AD models. (a) Schematic summarizing key MAPK signaling pathways including ERK1/2, p38, and JNK signaling cascades, the three families of MAPKs, all of which result in the activation of transcription factors which regulate gene regulation. (b) Experimental outline for Luminex studies of acutely-isolated CD11b+ CNS myeloid cells and frontal cortex brain samples isolated from age-matched WT and 5xFAD mice (n=7-9 mice/group). CD11b+ cells were isolated by Percoll density centrifugation followed by CD11b enrichment using MACS columns. (c-d) Summary of phospho-protein signaling data for ERK1/2, p38, and JNK signaling cascades in CD11b+ CNS myeloid cells (c) and frontal cortex samples (d) from WT and 5xFAD mice. Mean and SEM are shown.

To ascertain the expression levels of MAPKs at the total protein level in mouse microglia, we assessed the relative abundance of total MAPK proteins (as compared to non-MAPK proteins) in recently published proteomic datasets obtained from purified mouse microglia as well as BV2 mouse microglia. In total, 60 genes/proteins have been classified as belonging to the family of MAPK cascades (Supplemental Table 1) and this list of proteins was cross-referenced against two published microglial proteomes (Yates et al, 2017). 17 MAPK proteins (total protein levels) were identified in acutely isolated CD11b+ microglia from WT, 5xFAD and LPS-treated WT mice (Figure 2a), while 10 MAPK proteins were identified in BV2 microglia (Figure 2b). Interestingly, relative abundance of ERK proteins were highest in both acutely-isolated mouse microglia (Figure 2a), as well as in BV2 microglia (Figure 2b). Compared to WT microglia, 5xFAD microglia showed lower total protein levels of Raf1 and MAP2K6, while microglia isolated from LPS-treated mice showed increased total levels of two ERK proteins (MAPK1 and MAPK3) and lower levels of Taok1 and Raf1 (Figure 2c). Unlike our Luminex studies which showed alterations in levels of activated ERK phospho-proteins in 5xFAD microglia, total protein levels of ERK1 and ERK2 were not altered in 5xFAD microglia, indicating that activation of ERK signaling rather than increased protein expression is likely to occur in 5xFAD microglia. Consistent with this, immunofluorescence microscopy studies of wild-type and 5xFAD brains also confirmed no differences in total ERK protein levels (Figure 2d), while pERK1/2 expression was increased in microglia with activated morphology in 5xFAD brains (Figure 2e). In flow cytometric studies of acutely-isolated brain myeloid cells (Figure 2f), we also confirmed that intracellular pERK1/2 levels were increased in 5xFAD microglia as compared to wild-type microglia, whereas total ERK1/2 was equal between the groups (Figure 2f-g).

**Figure 2.**
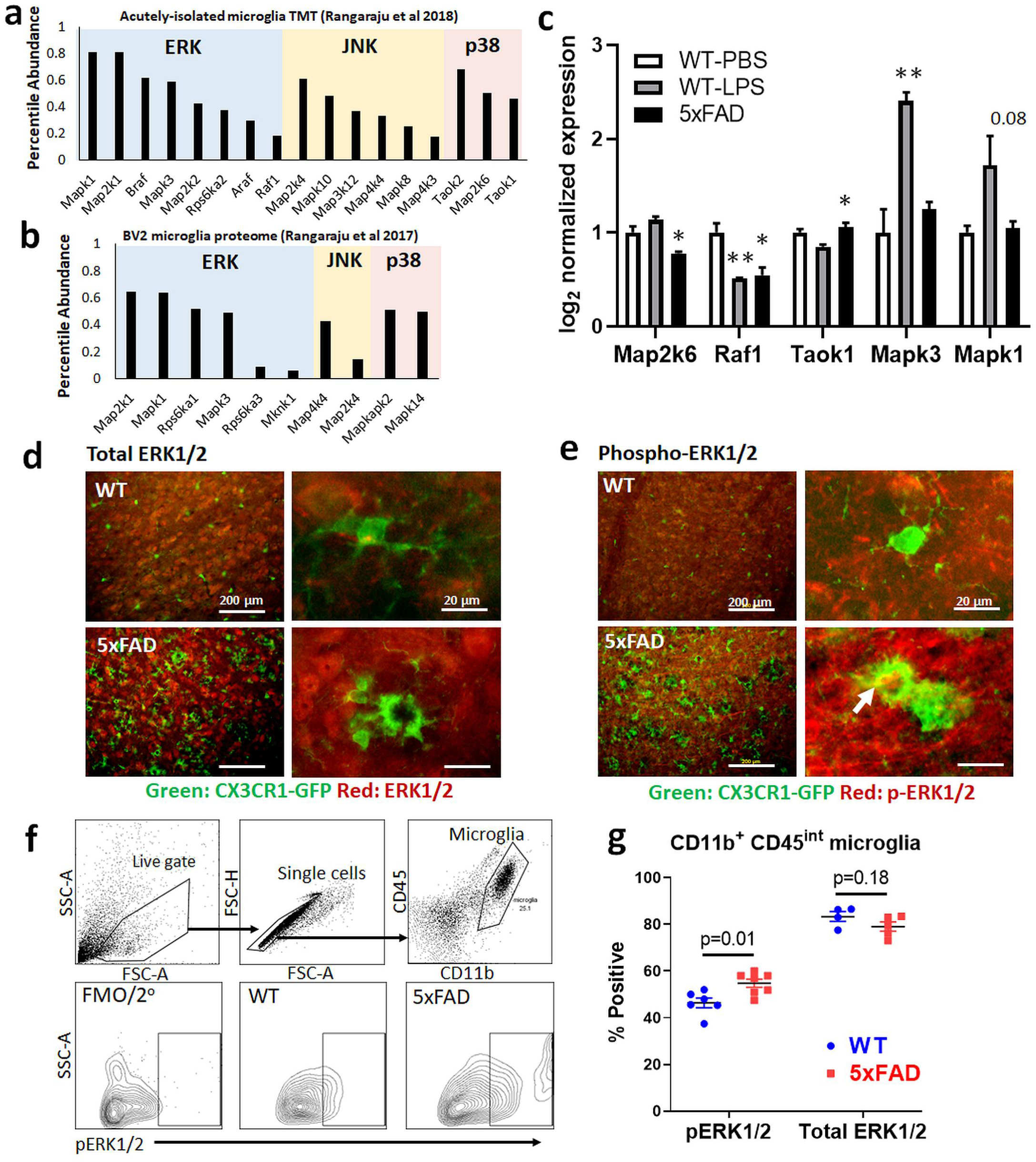
Confirmation of increased ERK1/2 activation in 5xFAD microglia. (a-b) Bar plot showing the relative abundance of 17 MAPK proteins as compared to non-MAPK proteins in microglia from WT, 5xFAD, and LPS-treated WT mice (a) and BV2 microglia (b) using published proteomic datasets. (c) Differentially expressed MAPK proteins identified in WT, 5xFAD, and LPS-treated WT mice. Error bars represent SEM. *p<0.05, **p<0.01, ***p<0.005. (d-e) Immunofluorescence microscopy of WT and 5xFAD brains probing for total ERK1/2 and p-ERK1/2 proteins in microglia. CX3CR1-GFP/WT and CX3CR1-GFP/5xFAD mice (age 7-8 mo) were used for these studies (n=3 mice/group). (f) Intra-cellular flow cytometric analysis of acutely-isolated CNS myeloid cells for p-ERK1/2 labeling. Gating strategy used is shown (top). (g) Quantitative analyses of flow cytometric studies of p-ERK1/2 and total ERK1/2 expression in WT and 5xFAD CD11b+ CD45int microglia.

In summary, our findings demonstrate that MAPK signaling proteins belonging to the ERK cascade are highly abundant in microglia and that microglia in AD mouse models exhibit up-regulation of ERK phospho-signaling, supporting critical roles for ERK signaling in regulating neuroinflammation in AD.

### ERK signaling is a critical regulator of pro-inflammatory activation in microglia

We recently reported that ERK signaling may be an important regulator of pro-inflammatory microglial activation. Based on our results showing increased ERK signaling in 5xFAD microglia and the relevance of ERK to human AD pathology, we next hypothesized that ERK is a key regulator of DAM gene expression, particularly those genes pertaining to pro-inflammatory responses that can be detrimental in the context of AD. To test this hypothesis, we exposed mouse primary microglial cultures isolated at postnatal day 0-3 to a selective ERK1/2 inhibitor (SCH772984, 1μM) in the presence or absence of IFNγ stimulation (1ng/mL) and performed viability studies as well as NanoString transcriptomic profiling for 770 neuroinflammatory genes. Overall, ERK inhibition minimally impacted cell health and viability under our experimental conditions (Supplemental Figure S1).

Of 770 genes in the NanoString panel, 678 were included for final analysis after normalization (Supplemental Table S2). One-way ANOVA identified 465 genes with differential expression across all 4 groups (436 genes also met FDR<0.05 criteria), and K-means clustering of these differentially expressed genes revealed 4 main clusters of genes (Figure 3a, Supplemental Table S3). Cluster 1 was comprised of genes negatively regulated by ERK under both unstimulated and IFNγ-stimulated conditions. Cluster 2-4 were all positively regulated by ERK signaling. Within these ERK-positively-regulated clusters, Cluster 2 was suppressed by IFNγ, Cluster 3 was upregulated by IFNγ and Cluster 4 was not affected by IFNγ. Among Cluster 3 genes, we also observed that the effect of ERK inhibition was most pronounced in the presence of IFNγ. Most statistically significant and representative genes for each cluster are shown in Figure 3a.

**Figure 3.**
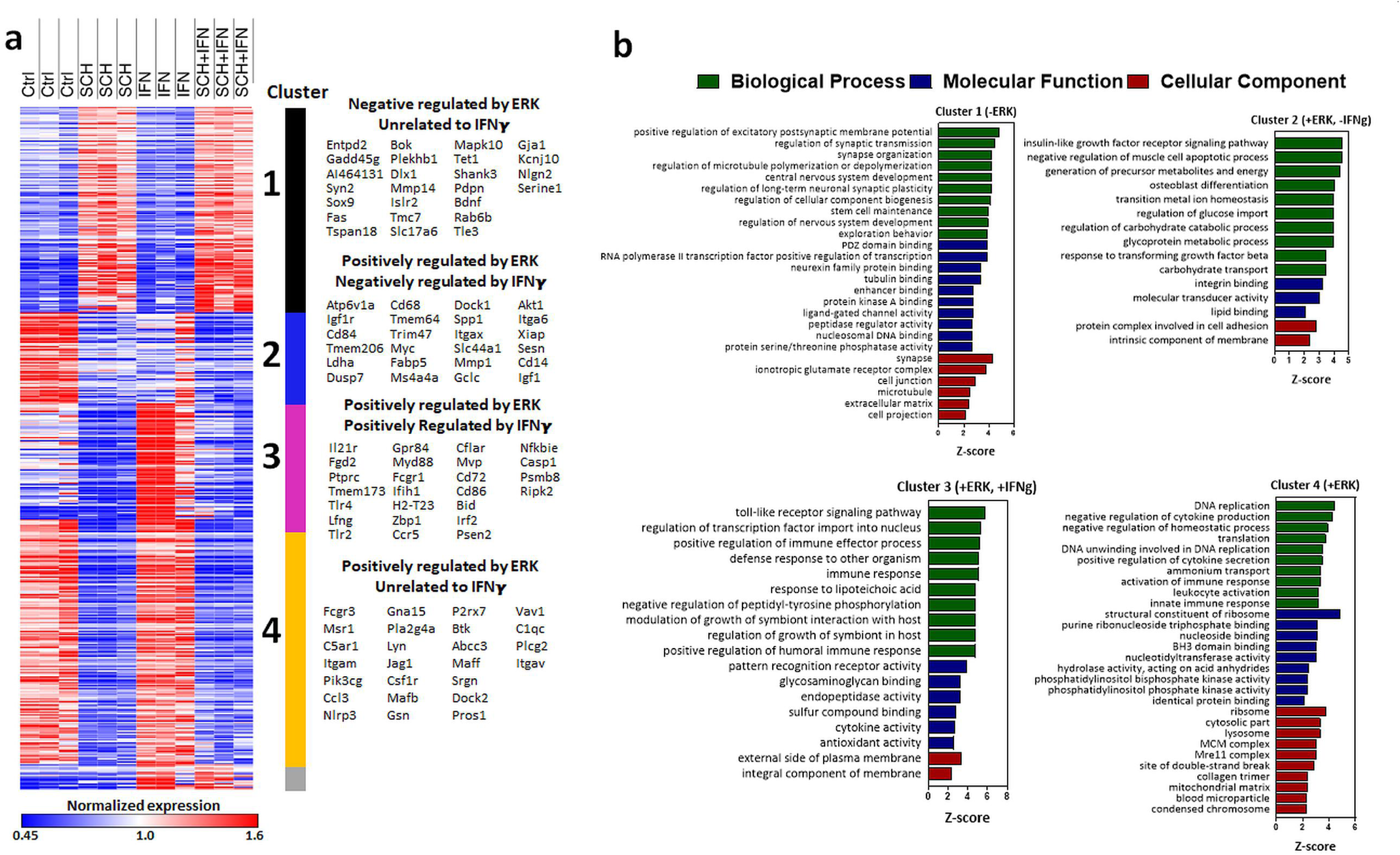
Transcriptomic profiling of microglia reveals distinct clusters of genes regulated by ERK1/2. (a) Heatmap showing K-means clustering analysis of 465 genes with differential expression across 4 treatment groups, as identified by one-way ANOVA, using NanoString gene expression data. Cluster 1: inhibited by ERK; Cluster 2: activated by ERK and inhibited by IFNγ; Cluster 3: activated by ERK and IFNγ; Cluster 4: inhibited by ERK. (b) Gene Ontology (GO) analysis depicting enrichment of GO terms in genes belonging to each cluster.

To obtain a broad functional understanding of the genes involved in each cluster, we next performed GO enrichment analyses (GOElite) on each cluster using all included genes in the neuroinflammation panel as the reference. Cluster 1 genes were enriched for neuronal/synaptic and glutamatergic transcripts with generally low abundance. Cluster 2 was enriched for genes involved in IGF1 receptor signaling, negative regulation of apoptosis, cell differentiation and energy metabolism and integrin binding. Cluster 3 was enriched with genes involved in pro-inflammatory signaling and immune processes, cytokine activity and transcriptional factor activity in the nucleus. Cluster 4 was enriched with genes involved in DNA replication, cell division, cytokine secretion, ribosomal machinery and purine metabolism (Figure 3b). Additional KEGG and Wikipathway enriched terms are shown in Supplemental Table S4. Since ERK signaling finally regulates transcription factor (TF) activity, we used bioinformatics approaches (Metacore, Thompson Reuters) to identify TFs down-stream of ERK activation that are likely to regulate gene expression in Cluster 2, 3 and 4 genes (Supplemental table S5). PU.1, c-Jun, Bcl-6, SP3, EGR1 and SP1 were top predicted TF regulators of Cluster 2 genes, consistent with known importance of PU.1, c-Jun and SP1 in microglial development and survival and the enrichment of homeostatic genes within cluster 2 (Rustenhoven et al, 2018; Smith et al, 2013). Similarly, Cluster 4 genes were also predicted to be regulated primarily by PU.1, Bcl-6, SP1 and c-Myc, again consistent with the enrichment of homeostatic and constitutive gene ontologies in this cluster. On the other hand, Stat1, NFκB, Bcl-6, IRF1 and PU.1 were predicted to regulate Cluster 3 genes that were enriched in pro-inflammatory genes. These agree with previous publications that demonstrated critical roles for Stat1, NFκB, IRF1 (along with IRF8) and PU.1 in microglial activation (Rustenhoven et al, 2018; Smith et al, 2013).

To determine the enrichment patterns of homeostatic, pro-inflammatory DAM and anti-inflammatory DAM gene expression within each cluster, we mapped our previously published lists of homeostatic, pro-inflammatory DAM, anti-inflammatory DAM as well as M1-like and M2-like gene lists (Rangaraju et al, 2018b) to a 2D tSNE representation of the gene clusters (Figure 4a-b). We observed that clusters 2 and 4 were enriched with homeostatic and anti-inflammatory DAM genes, while cluster 3 was enriched with pro-inflammatory DAM genes as well as M1-like genes (Figure 4b-c). In particular, ERK was found to positively regulate several canonical DAM genes, including Ccl3, Ch25h, Spp1, Igf1, and Itgax (Supplemental Figure S2) and pro-inflammatory DAM genes, including Ccl2, Irf1, Tnf, Slamf9, and Cd69 (Supplemental Figure S3). We also calculated a synthetic eigenvalue to represent overall expression of homeostatic, anti-inflammatory DAM and pro-inflammatory DAM profiles across each experimental condition using previously identified hub genes (Kme>0.75) of each microglial profile that were present in our Nanostring dataset (Dai et al, 2018; Gao et al, 2019). We confirmed that inhibition of ERK activity suppresses homeostatic, anti-inflammatory as well as pro-inflammatory DAM gene expression while IFNγ increases pro-inflammatory DAM and suppresses anti-inflammatory DAM gene expression (Figure 4d). A summary of the direction of regulation of each cluster by IFNγ and ERK activity is shown in Figure 4c. We also performed validation qRT-PCR studies using primary mouse microglia to confirm ERK-mediated regulation of several DAM genes (Apoe, Trem2, Tyrobp, Spp1) identified in Cluster 2 as well as pro-inflammatory genes (Tnf) identified in Cluster 3 (Supplemental Figure S4).

**Figure 4.**
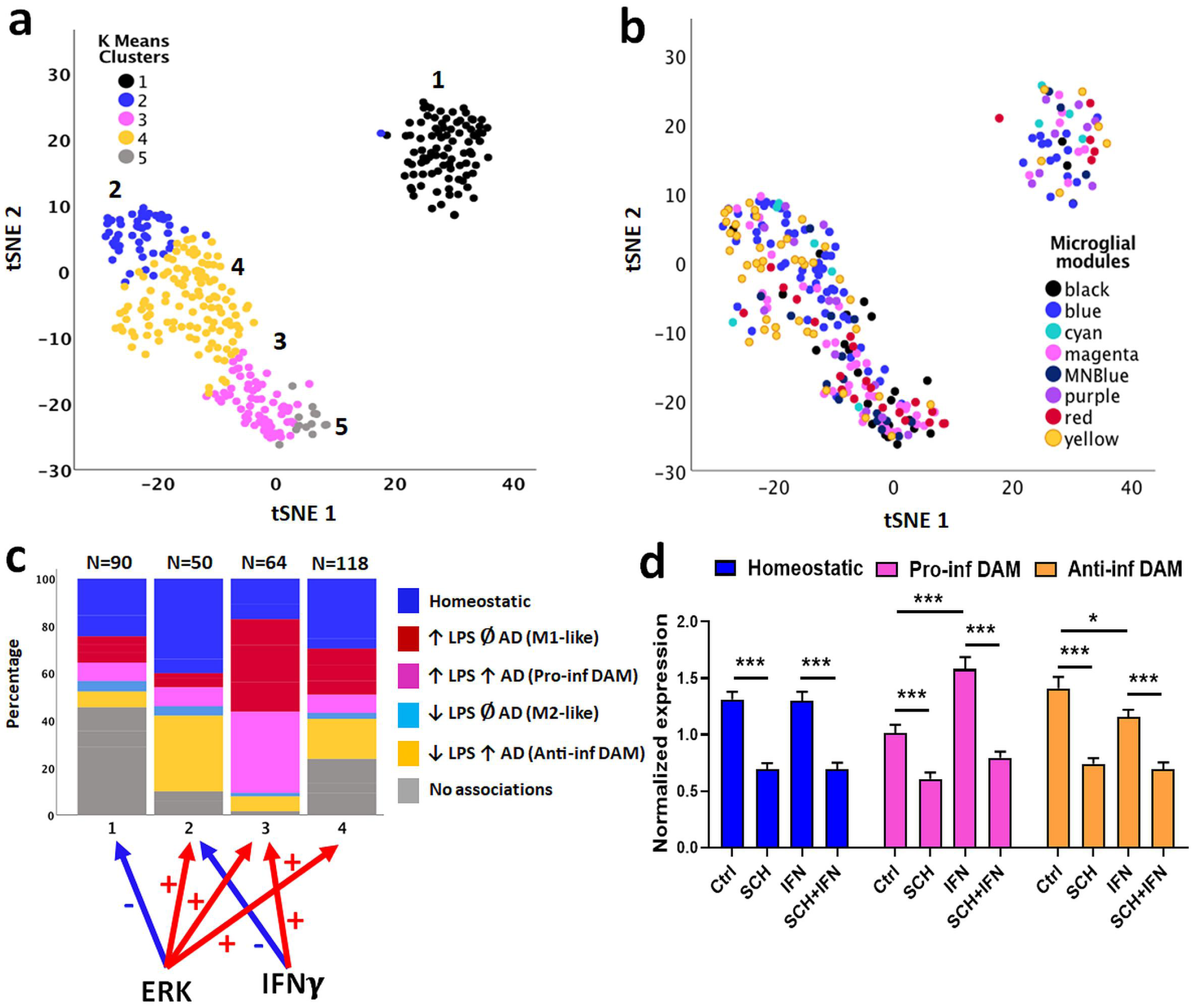
ERK1/2 positively regulates homeostatic, anti-inflammatory DAM, and pro-inflammatory DAM gene expression in microglia. (a) t-SNE plots showing clusters of genes based on NanoString expression data. Color coders indicate K-means clusters identified in Figure 3. (b) Mapping of previously identified microglia gene co-expression module hub genes across tSNE clusters. (c) Distribution of homeostatic, pro-inflammatory DAM, anti-inflammatory DAM, M1-like, and M2-like microglial genes in each cluster. (d) Expression of synthetic eigengenes of each microglial state (homeostatic, pro-inflammatory DAM and anti-inflammatory DAM) across experimental conditions. N=3 replicates per group. Error bars represent SEM. *p<0.05, **p<0.01, ***p<0.005.

To summarize our findings from transcriptomic and *in-vitro* microglial functional studies, ERK appears to be important for homeostatic, protective as well as pathological responses mediated by microglia. However, the critical role for ERK in regulating pro-inflammatory immune responses appears to be most evident under pro-inflammatory conditions. This suggests that the role for microglial ERK in promoting neuroinflammatory disease mechanisms may be highly context dependent. While ERK activation in homeostatic microglia may regulate constitutive functions and mediate the effects of several trophic factors, ERK activation under disease-associated conditions, such as in AD, may regulate pro-inflammatory immune responses.

### ERK signaling regulates microglial phagocytic function

Having found that ERK signaling regulated diverse DAM-associated genes, we next asked whether or not ERK inhibition would modulate microglial phagocytic function. To test this, we used a flow cytometric assay of fluorescently labeled Aβ42 fibril phagocytosis. We observed that ERK inhibition reduced the ability of microglia to phagocytose Aβ42 fibrils regardless of stimulation by IFNγ (Figure 5a-b). Since pro-inflammatory microglia can also mediate neurotoxicity via direct neuronal phagocytosis, we performed *in-vitro* microglia/neuronal co-culture studies and determined the effect of IFNγ and ERK inhibition on microglia-mediated neuronal phagocytosis. N2a cells were stably transduced to express GFP and then differentiated with retinoic acid under low serum conditions for 4 days. Primary mouse microglia, which were treated with IFNγ and/or ERK inhibitor were then added to differentiated N2a-GFP cells for another 24 h, after which microglia were collected for flow cytometric studies to measure GFP positivity within microglia. First, we observed that addition of microglia, regardless of treatment, resulted in decreased viability in N2a cells, indicating that *in-vitro* cultured primary microglia induce some level of neurotoxicity, possibly due to baseline level of activation as a result of *in-vitro* culture conditions and serum exposure (Figure 5c). Inhibition of ERK significantly reduced microglial phagocytosis of N2A cells with or without IFNγ treatment (Figure 5d). These functional assays support a role for ERK in phagocytic activity, both for Aβ as well as neuronal phagocytosis.

**Figure 5.**
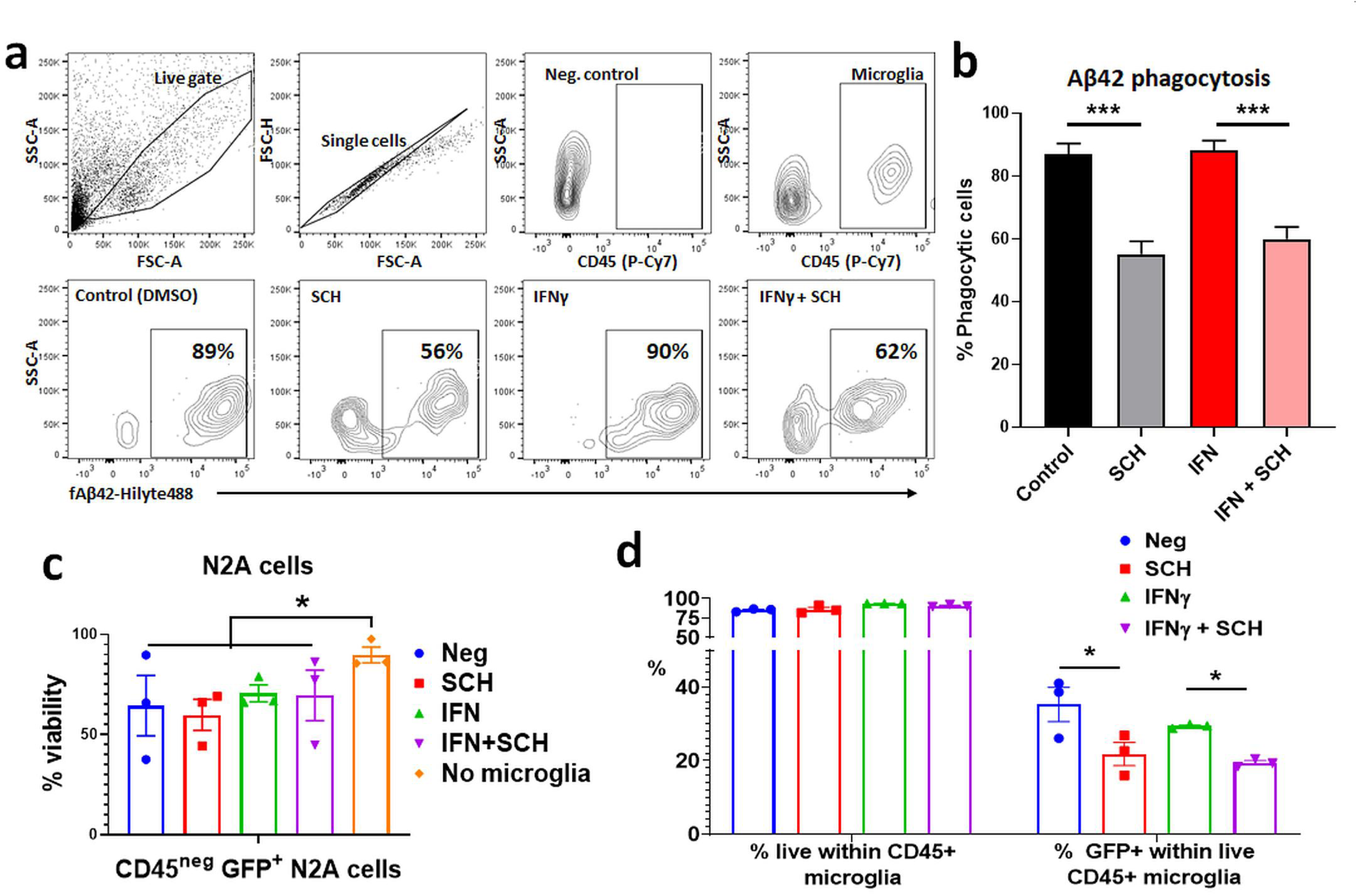
Roles of ERK activity in regulating microglial phagocytosis of fibrillar Aβ42 and of neuronal phagocytosis. (a) Flow cytometry experimental data showing the uptake of fluorescent Aβ42 fibrils by live CD45+ microglia that had been treated with an ERK1/2 inhibitor and/or IFNγ. (b) Bar graphs showing the reduction of microglial phagocytosis of Aβ42 by ERK inhibition under unstimulated and stimulated conditions *in-vitro*. N=3 replicates per group. Error bars represent SEM. *p<0.05, **p<0.01, ***p<0.005. (c) Bar graphs showing a reduction of microglial phagocytosis of differentiated N2a cells in ERK inhibited groups. GFP+ N2a cells were differentiated and then co-cultured with primary microglia that were pre-treated with an ERK inhibitor with or without IFNγ. N=3 replicates per group. Error bars represent SEM. *p<0.05, **p<0.01, ***p<0.005. (d) Bar plot demonstrating a reduction in the viability of N2a cells when co-cultured with microglia. N=3 replicates per group. Error bars represent SEM. *p<0.05, **p<0.01, ***p<0.005.

### A framework for ERK activation down-stream of receptor tyrosine kinases (RTKs) in homeostatic microglia and DAM

Since ERK signaling is initiated by activation of receptor tyrosine kinases (RTKs) on the cell surface, it is likely that the predominant types of RTKs expressed by cells, are the key determinants of ERK activation (Figure 6a). ERK activation via one RTK can have very different effects on cellular gene expression as compared to another RTK. This variability is attributed to differences in the kinetics of ERK activation, as well as simultaneous activation of other RTK-mediated signaling pathways such as PLC-mTOR, other MAPK signaling, NFκB and others. Microglia express several types of RTKs, each with distinct ligands and cellular functions. Based on our recently published transcriptomic landscape of distinct microglial activation states (Rangaraju et al, 2018b), we shortlisted RTK genes that were highly co-expressed with homeostatic, M1-like, M2-like, anti-inflammatory DAM as well as pro-inflammatory DAM gene profiles. This analysis revealed that homeostatic microglia highly express two RTKs, Csf1r and Mertk. The critical role for Csf1r in regulating microglial survival in mice has been well established. Csf1r is exclusively expressed by microglia in the brain and its inhibition leads to rapid depletion of homeostatic microglia in mouse models (Konno et al, 2018; Spangenberg et al, 2019). Moreover, Csf1r mutations in humans are associated with neurodegenerative disease conditions as well as abnormalities in brain development (Guo et al, 2019; Oosterhof et al, 2019). Mertk is a member of the TAM (Tyro3, Axl and Mertk) RTK family, is involved in phagocytic activity, and is primarily expressed by astrocytes and microglia in the brains of both humans and mice (Zhang et al, 2014). In DAM gene clusters, we found that Axl (another TAM member like Mertk) and Flt1 were characteristic of and elevated in expression in the canonical DAM state. Axl has been consistently identified as being highly expressed by DAM or neurodegeneration-associated microglial states (Fourgeaud et al, 2016; Yin et al, 2017). Within DAM sub-profiles, interestingly, Flt4 was highly co-expressed in the pro-inflammatory (Magenta) DAM gene cluster while Igf1r was highly co-expressed in the anti-inflammatory (Yellow) DAM gene cluster (Figure 6b). Given this distinct pattern of RTK expression by homeostatic microglia and DAM, we searched the literature for known ligands specific for each RTK. These ligands have been shown in Figure 6c. This framework suggests that ERK signaling in homeostatic microglia is likely to be primarily driven by IL34 and CSF1, both ligands of Csf1r, as well as agonists of Mertk such as Galectin3 and Gas6. On the other hand, ERK signaling in DAM may be primarily regulated by Axl and Flt1, and the pro-inflammatory and anti-inflammatory sub-profiles within DAM likely may utilize Flt4 and Igf1r to further fine tune ERK signaling, respectively. This hypothetical framework for RTKs and RTK-mediated ERK signaling in homeostatic and DAM profiles in AD can guide future investigations to identify approaches to regulate ERK activity in microglia in order to obtain a desired neuroinflammatory phenotype in neurodegenerative disorders.

**Figure 6.**
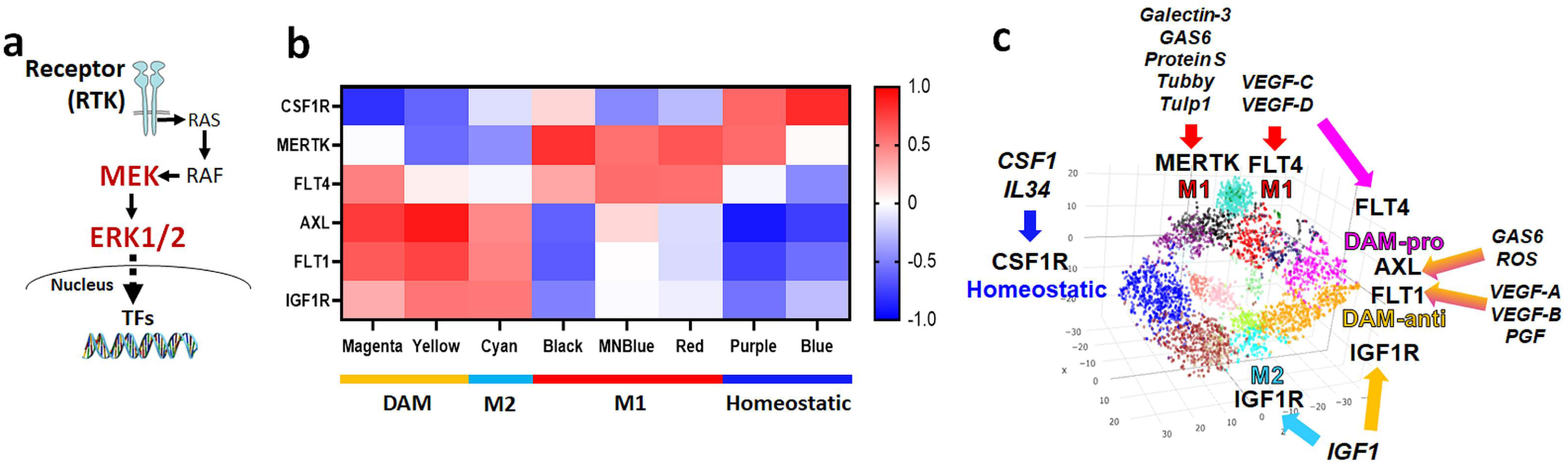
Identification of receptor tyrosine kinases (RTKs) that signal via ERK in homeostatic microglia and DAM states. (a) Schematic depicting axis of signal transduction for the MAPK/ERK pathway, beginning with RTK signaling and resulting in the activation of transcription factors. (b) Heatmap displaying K-means clustering of 6 RTK genes that mapped to DAM, M2-like, M1-like, and homeostatic modules in previously published analyses of microglia transcriptomes. (c) Transcriptomic framework of microglial activation showing RTK expression relative to each module and the ligands that activate each of the RTK’s.

### GWAS and proteomic datasets reveal evidence for a pathophysiological role for ERK activation in human AD

If ERK signaling in microglia is important in human AD pathogenesis, we predicted that ERK signaling should lie upstream of immune determinants of AD genetic risk. To test this hypothesis, we cross-referenced ERK regulated gene clusters against late-onset AD risk genes identified in meta-analyses of human GWAS studies, using an approach called MAGMA (de Leeuw et al, 2015; Seyfried et al, 2017). Of over 1,000 known AD risk genes, 43 were present in the NanoString panel and 28 demonstrated differential expression in our dataset, mapping to any one of the 5 gene clusters (Figure 7a, Supplemental Table S6). While there was no statistically significant over-representation of AD risk genes in any of the clusters, risk genes with highest MAGMA significance mapped to Cluster 2 (Cnn2, Ms4a4a, Mef2c, Cd33) and Cluster 4 (Bin1, Pilra, Ep300). Cluster 3 AD risk genes included Sqstm1, Lrrc25 and Nrp2. Of these 28 ERK-regulated AD risk genes, 11 genes are specifically expressed in microglia (highlighted in Figure 7a with red asterisks).

**Figure 7.**
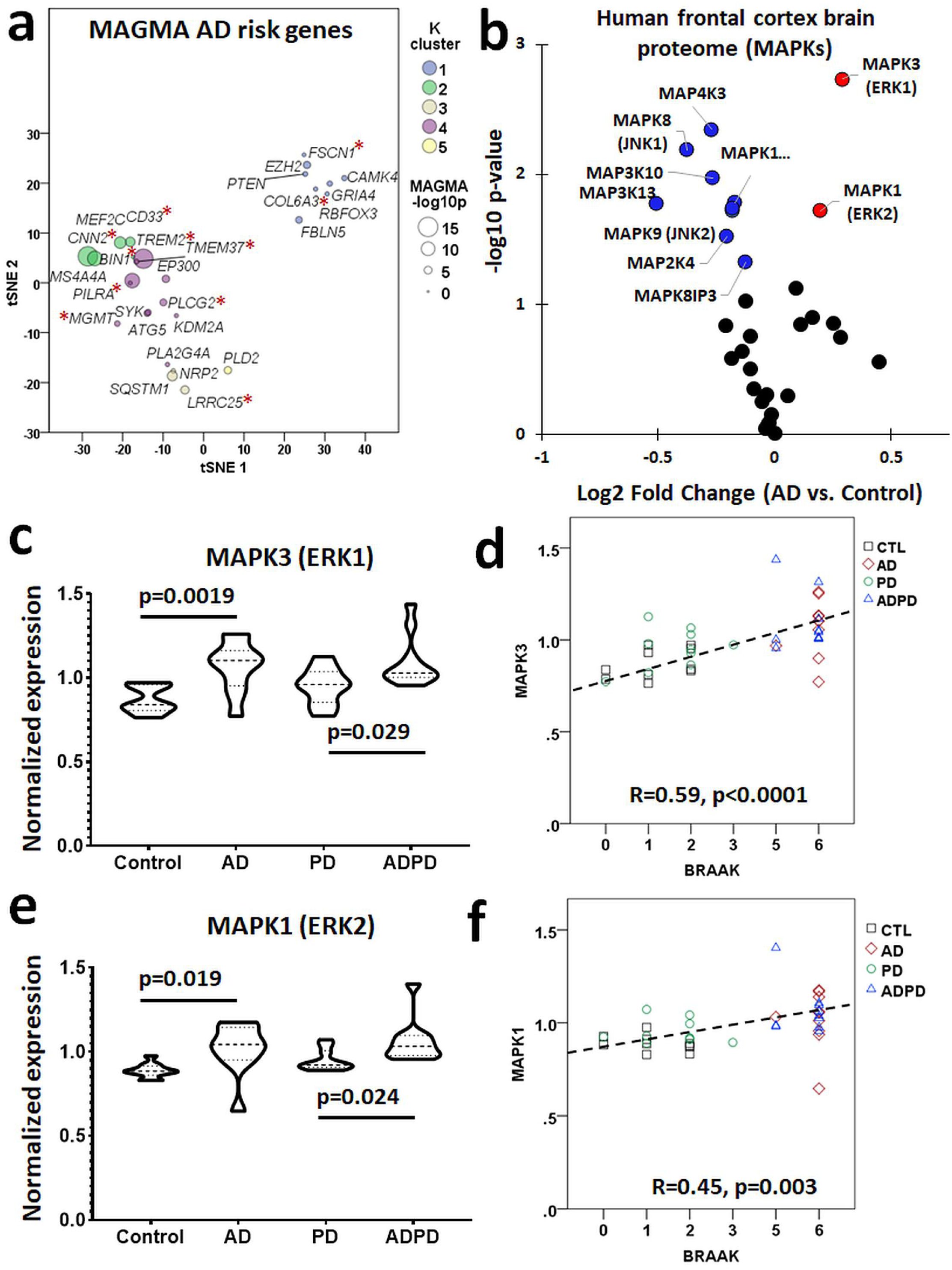
MAGMA and TMT proteomics reveals role for ERK activation in human AD. (a) tSNE plot showing late-onset AD risk genes identified in human GWAS analyses that are present in the NanoString dataset (*p <□.05). (b) Differential expression analysis of human frontal cortex proteomes showing that MAPK1 (ERK2) and MAPK3 (ERK1), both shown in red, are expressed more highly in AD brains than in control brains while the opposite trend is observed in other notable MAPKs, shown in blue. (c) Violin plots showing that MAPK3 expression levels are increased in AD and ADPD brains as compared to PD or control brains (n=10 per group). (d) Linear regression depicting a positive correlation between BRAAK stage and MAPK3 expression. (e) Violin plots showing that MAPK1 expression levels are increased in brains with AD pathology (n=10 per group). (f) Linear regression depicting a positive correlation between BRAAK stage and MAPK1 expression.

To determine whether MAPK pathway protein expression and phospho-signaling is altered in human AD, we investigated two post-mortem brain quantitative proteomic datasets. The first dataset was derived from 10 controls, 10 AD cases, 10 with Parkinson’s disease and 10 with both AD and PD pathological features. Tandem mass tag proteomics was performed on brain tissues obtained from the dorsolateral pre-frontal cortex (Supplemental Table S7). In this dataset, 33 MAPK proteins were identified and differential expression analyses comparing AD with control samples revealed increased expression of MAPK1 (ERK2) and MAPK3 (ERK1) in AD, while other MAPKs including MAPK8-10 (JNK1-3) were decreased in AD brains (Figure 7b). We also found that increased levels of MAPK1 (ERK2) and MAPK3 (ERK1) were only seen in brains with AD pathology while PD brains did not exhibit increased MAPK1 or MAPK3 levels (Figure 7c-e). We also observed strong associations between AD neuropathological grade (BRAAK stage) and protein levels of MAPK1 (R=0.45, p<0.005) and MAPK3 (R=0.59, p<0.005) (Figure 7d,f). These findings from total protein expression data show that ERK signaling proteins are specifically increased in AD brains. However, these data do now confirm whether activation of ERK proteins are also increased.

We then turned to a phospho-proteomic dataset in which frontal cortex samples from 6 controls, 6 asymptomatic AD pathology and 6 symptomatic AD cases were used for quantitative TMT proteomics after purification of phosphopeptides on an immobilized metal affinity chromatography (IMAC) column. A comprehensive description and analysis of this phosphoproteomic dataset will be published separately and the expression data and methods have been deposited online at http://proteomecentral.proteomeexchange.org. Within this dataset, we obtained expression data for phosphopeptides that mapped to any of the known 60 MAPK family reference protein/gene list. This identified 127 phospho-peptides of which 57 peptides showed differential expression (unadjusted ANOVA p<0.05) across controls, asymptomatic AD and symptomatic AD cases (Figure 8, Supplemental Table S8). K-means clustering of these differentially expressed peptides revealed distinct clusters of peptides with unique trajectories of change across stages of AD progression (control→AsymAD→AD) (Figure 8a). Cluster 1 comprised of peptides with dose-dependent increase across the three groups, and Cluster 4 showed early and sustained increase, while Cluster 3 showed a delayed increase. Conversely, Clusters 2 showed decreased expression in clinical stages of AD pathology although the number of these peptides were relatively small. The overall trajectories of change for each cluster are presented in Figure 8b. Based on known biological functions of each MAPK protein, we found that 20 peptides (11 gene symbols) mapped to the ERK cascade, while 23 peptides (8 gene symbols) mapped to JNK, 8 peptides (3 gene symbols) mapped to p38, and 4 peptides mapped to MAPK proteins involved in cross talk between JNK and p38 MAPK cascades (Figure 8c). To gain further insights into roles of differentially expressed phosphopeptides in each signaling pathway, we then mapped each phosphopeptide to known MAPK signaling pathways. ERK-related peptides mapped to all levels of the ERK signaling cascade (RAF→MEK→MAPK→RSK) (Figure 8d). Within the 20 ERK-related phosphopeptides with differential expression in this dataset, we then identified key phosphosites in ERK-family proteins of AD relevance. Three RSK peptides (RPS6KA2 pS434 and pS402, and RPS6KA5 pS750) and RAF peptides (BRAF dual phosphorylated pS358+pS750 and ARAF pS157) were increased at least 1.5-fold in AD compared to controls, and 2 RSK phosphopeptides (RPS6KA2 pS434 and pS402) were also increased 1.5-fold in the asymptomatic stage of AD (Supplemental Table S8). Two phosphosites in ERK2 were also increased in AD (MAPK1/ERK2 pS284, MAPK1/ERK2 pS202). While little is known about pS284 and its impact on ERK activity, S202 in ERK2 is located at the activating lip in very close proximity to two phosphorylation sites that confer full ERK kinase activity (ERK2 T185/Y187), and a S202P mutation confers resistance to ERK inhibitors, suggesting that ERK phosphorylation at S202 may positively regulate ERK activation (Goetz et al, 2014). In addition to ERK phosphopeptides, we also found that early signaling events in the JNK and p38 cascades represented by phosphopeptides were also differentially expressed in AD brains (Supplemental table S8). To identify upstream triggers that may regulate these differentially expressed MAPK phosphopeptides in AD, we performed pathway enrichment analysis using PANTHER and found that the majority of proteins were downstream of growth factor receptors and receptor tyrosine kinases (RTKs) such as PDGF, GnRH, FGF, EGF, TGF and VEGF receptors (Supplemental table S9). These findings using cross-sectional human post-mortem brain data show unique patterns of ERK activation in asymptomatic and symptomatic stages of AD pathology, suggesting that differential flux through MAPK signaling pathways, specifically ERK, may underlie symptomatic progression and neurodegeneration. We have summarized these patterns of differential ERK signaling observed in our phoshoproteomic analyses, in Figure 9. Overall, ERK signaling activation in human AD brains was consistent with our observations in the 5xFAD mouse model of Aβ pathology of AD.

**Figure 8.**
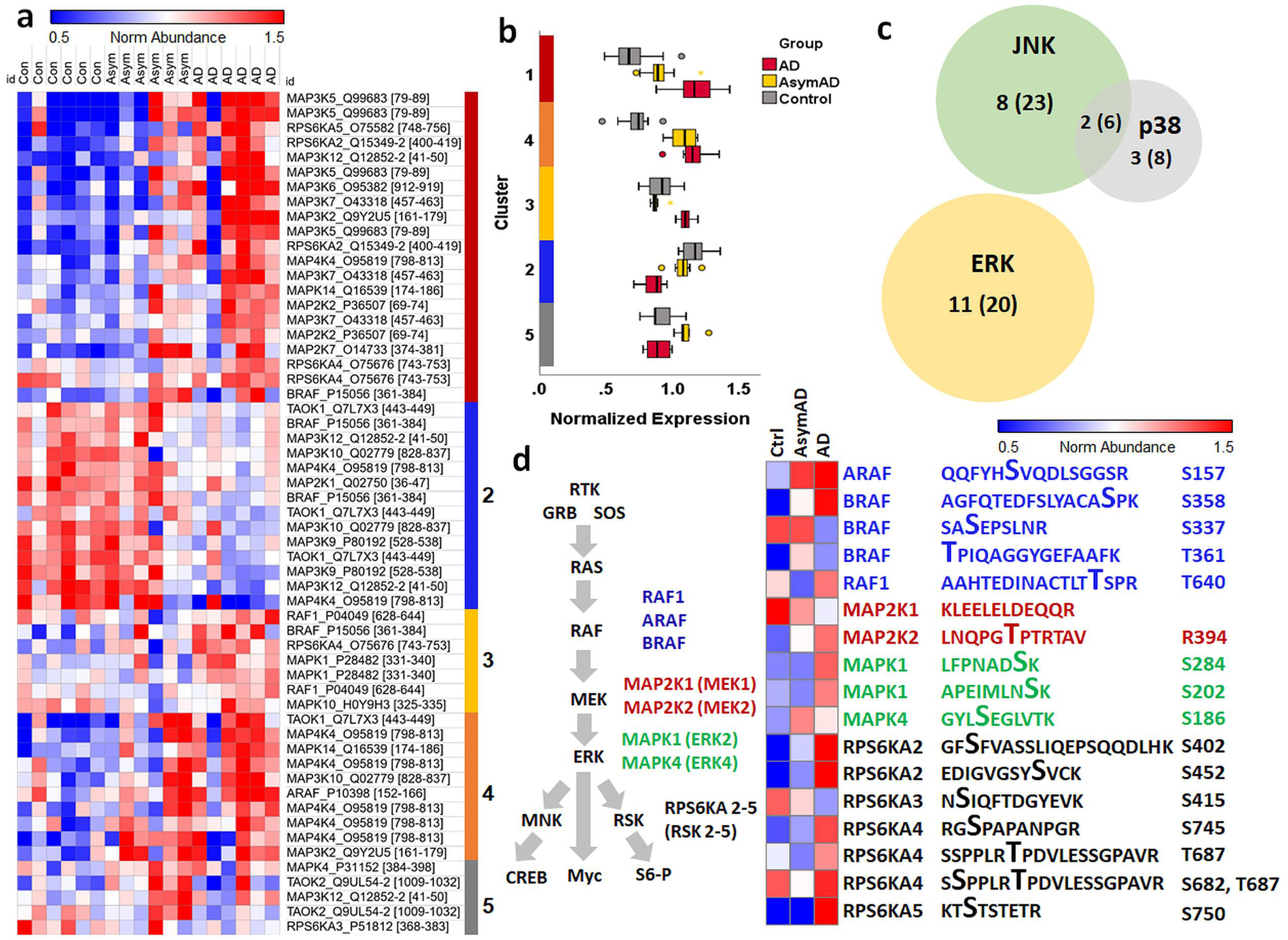
Phospho-proteomic analysis of human post-mortem brain reveals ERK activation in AD. (a) Heat map showing k-means clustering of 57 differentially expressed phospho-peptides that mapped to a MAPK reference gene list, across frontal cortex samples from 6 non-AD control, 6 asymptomatic AD, and 6 symptomatic AD cases. (b) Box and whisker plot quantifying the trajectory of change in expression for each cluster across stages of AD progression. (c) Diagram showing the number of peptides (in parenthesis) and their respective gene symbols (outside of parenthesis) are involved in ERK, JNK, and p38 signaling. (d) Visual representation and k-means clustering of key phosphosites within the differentially expressed ERK-related peptides in the IMAC dataset, mapping to all levels of the ERK cascade, that are relevant in AD.

**Figure 9.**
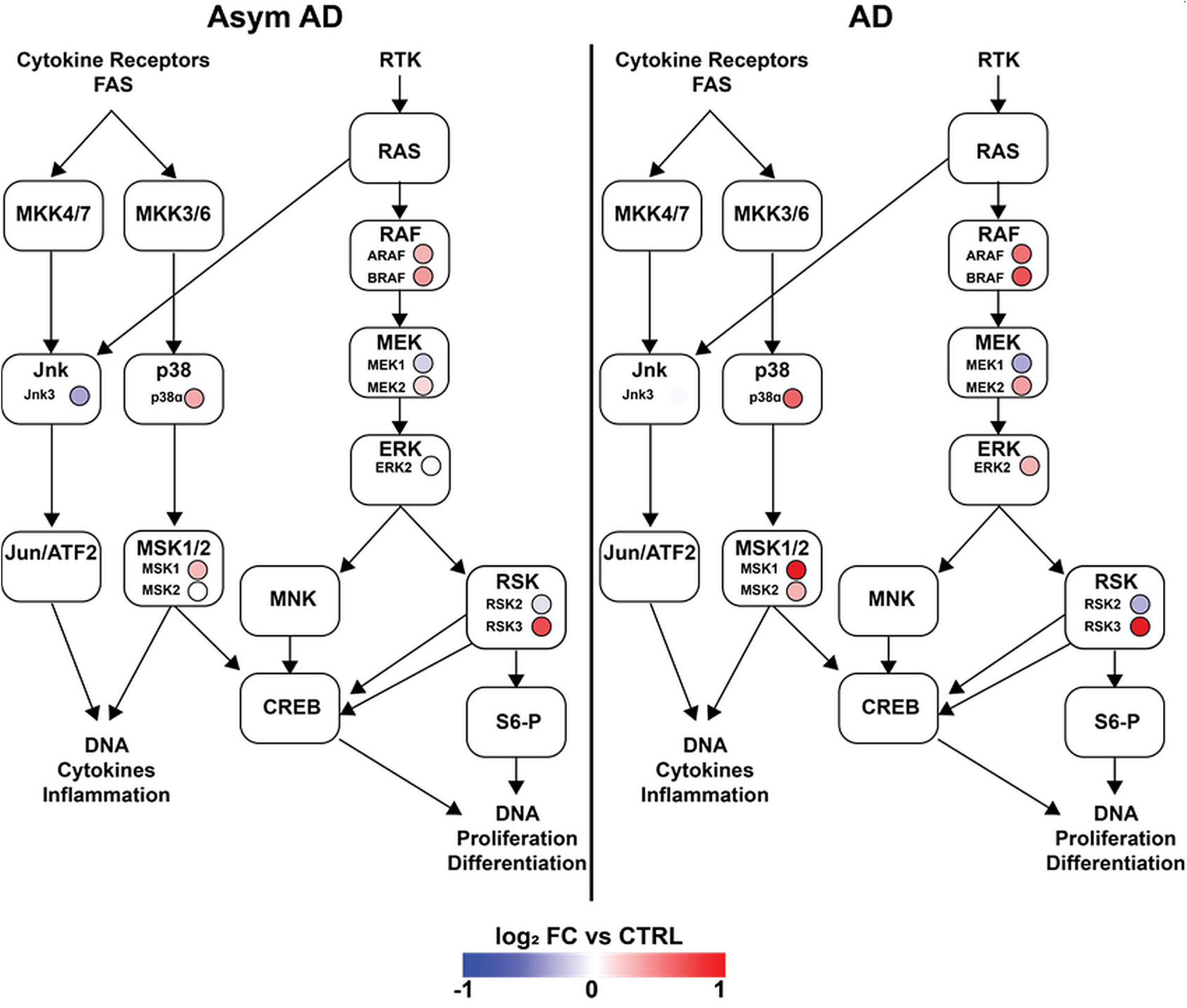
Phospho-proteomics analysis reveals progressively up-regulated flux through the MAPK pathway in asymptomatic AD and cognitively impaired AD cases. Colored circles represent fold-change of each detected phospho-protein protein compared to control cases (N=6 non-AD control, 6 asymptomatic AD, and 6 symptomatic AD cases).

## DISCUSSION

Immune responses are orchestrated by early phospho-signaling events that follow activation of cell surface receptors. In turn, phospho-signaling results in regulated expression of groups of genes that together determine the immunological phenotype and function of the cell. Among the various immune signaling cascades that may regulate microglial functions, we focused on the MAPK family of signaling proteins in the 5xFAD pre-clinical AD model in order to identify pathologically relevant changes in MAPK pathway signaling in microglia that is responsible for pro-inflammatory immune activation in AD. Using Luminex analyses of phospho-proteins in acutely-isolated microglia, we found that among the three main classes of MAPKs, up-regulated ERK signaling was the most apparent in microglia isolated from mouse AD models. However, no significant changes in MAPK activation were observed in the whole brain, indicating that up-regulated ERK activation is particularly characteristic of microglia in AD. Since ERK signaling is often transient following immune stimuli, our observation of significantly increased pERK1/2 levels in mouse microglia from AD models also suggests that ERK activation may be persistently elevated in AD, potentially as a result of sustained activation of ERK via agonists of RTKs.

In support of a disease-relevant role for ERK in AD, we also found that ERK proteins are highly abundant in microglia isolated from 5xFAD mice. Moreover, *in-vitro* experiments using primary microglia, revealed that inhibition of ERK strongly suppressed pro-inflammatory immune activation induced by IFNγ, which is an AD relevant pro-inflammatory cytokine (Wood et al, 2015; Zheng et al, 2016). We also found that ERK signaling is important for homeostatic and constitutive microglial functions including phagocytosis of Aβ, and that it was required for neuronal phagocytosis, which is a potentially neurotoxic microglial function duration AD pathogenesis (Rajendran & Paolicelli, 2018). ERK signaling was also found to be an upstream regulator of expression of several canonical DAM genes including Trem2 and Tyrobp, suggesting that ERK signaling may regulate the transition of homeostatic microglia to DAM, a phenotype commonly seen in AD and other neurodegenerative disease conditions. However, we did not observe any role for ERK signaling in regulation of microglial Apoe expression, suggesting that the Apoe-Trem2-dependent checkpoint for homeostatic-to-DAM transition in AD (Krasemann et al, 2017) may be regulated by both ERK-independent and ERK-dependent mechanisms. Interestingly, ERK activation was also upstream of several immune AD risk genes, including Cd33, Bin1, Plcg2, Trem2 and Cnn2, emphasizing the importance of ERK signaling in regulating immune pathways responsible for AD pathogenesis.

Consistent with these important roles for ERK signaling in mouse microglia and in 5xFAD AD mice, we found that human AD is also characterized by increased ERK activation. Our analyses of human proteomics data, including phospho-proteomics studies, confirmed that ERK1 and 2 are the only two MAPKs that are increased at the total protein level in human AD brains, while other MAPKs such as JNKs are decreased. At the phospho-protein level, 57 MAPK phosphopeptides were found to be differentially expressed in human AD brains, and the relative proportion of ERK proteins with differentially expressed phosphopeptides was highest compared to other MAPK proteins. Overall, our integrated analyses of signaling data obtained from pre-clinical 5xFAD AD mice and *in-vitro* mechanistic studies in microglia along with confirmatory human post-mortem brain proteomics findings, strongly support an important role for ERK signaling in microglia in AD pathogenesis.

From a therapeutic perspective, our results suggest that inhibition of ERK in microglia should limit pro-inflammatory disease mechanisms. However, our results also clearly highlight the importance of ERK in homeostasis and constitutive cellular functions, such as cell division, survival and phagocytosis. Furthermore, ERK is ubiquitously expressed in the brain in several cell types and is also important in neuronal physiological functions, including learning and memory (Hutton et al, 2017; Silingardi et al, 2011). Therefore, therapeutic inhibition of ERK signaling specifically in microglia will likely be limited by cellular non-selectivity, potentially resulting in undesired side effects. Rather than global inhibition of ERK activation, another potential strategy to attain desirable neuro-immunomodulatory effects would be to modulate microglial RTKs that primarily signal via ERK. We found that homeostatic and DAM, and pro- and anti-inflammatory DAM sub-profiles, are characterized by expression of distinct RTKs (Figure 6). While homeostatic microglial ERK signaling is likely to be regulated by CSF1R and MERTK, DAM may generally utilize AXL and FLT1. Inhibition of CSF1R is a well-known strategy to selectively and effectively deplete microglia in mouse models, emphasizing the critical role that this RTK and downstream ERK signaling are likely to play in survival of microglia. TAM receptors including MERTK and AXL have also been shown to be critical for microglia-mediated phagocytosis and clearance of debris (Fourgeaud et al, 2016). Within DAM sub-profiles, our results suggest that co-expression of FLT4 by pro-inflammatory DAM may regulate pro-inflammatory responses while IGF1R expression by anti-inflammatory DAM, may confer a protective phenotype. If distinct RTKs activate ERK in distinct manners, each resulting in expression of distinct gene sets, the dynamics of ERK signaling downstream of RTKs are likely to modulate downstream gene expression, although this has yet to be explored in microglia. A dynamic model of ERK activation in microglia raises the intriguing possibility that short-term ERK activation regulates homeostatic and constitutive responses while sustained and long-term ERK activation may mediate disease-associated and pro-inflammatory microglial responses in neurodegeneration. Indeed, *in-vitro* and *in-vivo* studies using real-time ERK signaling reporters in neuronal cells and T cells have shown that the immune effects of short-term ERK activation and sustained ERK activation are very different (de la Cova et al, 2017; Harvey et al, 2008; Konishi et al, 2018). The recent development of FRET-based biosensors of ERK and MAPK signaling will allow us to test mechanistic hypotheses to define the ERK signaling dynamics of diverse RTKs expressed by microglia, and then further determine the downstream effects of ERK activation via RTKs on gene expression patterns both *in-vitro* and *in-vivo*. These novel approaches will also facilitate the development of RTK modulators that regulate ERK signaling dynamics to achieve desired neuroinflammatory microglial phenotypes.

Although we focused on ERK signaling in microglia, our Luminex studies also suggested increased activation of p38 MAPK as well as trends towards increased JNK activation in microglia from 5xFAD mice and human post-mortem brain proteomes also suggested activation of JNK and p38 MAPK pathways, although to a lesser extent than ERK. These data suggest that non-ERK MAPKs may also be relevant in regulating microglial functions in AD. Beyond these MAPKs, NFκB signaling is another key regulator of pro-inflammatory immune responses, which was not considered in our studies (Shih et al, 2015). Unlike the robust effects of ERK inhibition observed in these studies, we previously found that inhibition of JNK or NFκB in microglia had minimal effects on the expression of selected homeostatic and DAM genes or on effects of pro-inflammatory stimuli in microglia (Gao et al, 2019). Due to the limited yield of protein from acutely-isolated CD11b+ microglia from adult mice, we focused our efforts on ERK, although future studies will investigate the importance of other MAPKs, specifically p38 and JNK signaling pathways, in AD.

In conclusion, our integrated analysis of a pre-clinical model of AD pathology, *in-vitro* studies in primary microglia, and human proteomic data provide novel insights into ERK activation in microglia as a critical regulator of pro-inflammatory immune responses and a potential immune mechanisms of AD pathogenesis. However, ERK activation in microglia is also necessary for homeostatic functions, indicating that global ERK inhibition may not be an appropriate immunomodulatory strategy in AD. Based on our framework of expression of distinct receptor tyrosine kinases by homeostatic and disease-associated microglia in AD, we propose that beneficial regulation of ERK signaling in AD may be achieved by modulating upstream receptor tyrosine kinases.

## ACKNOWLEDGEMENTS

This study was supported by the Alzheimer’s Association AARG 37102 (PI: Rangaraju), NIH K08-NS099474-1 (PI: Rangaraju), the Accelerating Medicine Partnership for AD (U01 AG046161 and U01 AG061357, PI: Levey), the Emory Alzheimer’s Disease Research Center (P50 AG025688, PI: Levey) and P30 AG066511(PI: Levey), and by startup funds from the George W. Woodruff School of Mechanical Engineering at the Georgia Institute of Technology (Wood). L.D.W. was supported in part by the National Institutes of Health Cell and Tissue Engineering Biotechnology Training Grant (T32-GM008433). This study was supported in part by the Emory Flow Cytometry Core (EFCC), one of the Emory Integrated Core Facilities (EICF) and is subsidized by the Emory University School of Medicine. Additional support was provided by the Georgia Clinical & Translational Science Alliance of the NIH under Award Number UL1TR002378. The content is solely the responsibility of the authors and does not necessarily reflect the official views of the NIH.

## AUTHOR CONTRIBUTIONS

MJC, SRamesha, LW and SRangaraju designed the study, and analyzed and interpreted the results. MJC, SRamesha, LDW, LW, TG, HX, LP, EBD and DD conducted the experiments. AIL, NTS, JJL and LP provided the human brain proteomic data for analysis. MJC, SRangaraju and LW wrote the article. All authors contributed to the article. SRangaraju and LW supervised the project.

## SUPPLEMENTAL INFORMATION

This manuscript contains 4 Supplemental Figures

Supplemental Figure 1. ERK inhibition exerts minimal impact on viability of microglia.

Supplemental Figure 2. DAM genes positively regulated by ERK signaling.

Supplemental Figure 3. Pro-inflammatory DAM genes positively regulated by ERK signaling

Supplemental Figure S4. qRT-PCR validation confirms ERK effect of select DAM genes.

This manuscript contains 9 Supplemental Tables

Supplemental Table S1. List of MAPK genes/proteins cross-referenced with published proteomic data sets.

Supplemental Table S2. NanoString transcriptomic dataset for 678 genes with ANOVA p-values.

Supplemental Table S3. K-means clustering of 465 differentially expressed genes in NanoString dataset.

Supplemental Table S4. Gene Ontology analysis of K-means clusters showing enriched GO terms, KEGG, and Wikipathways.

Supplemental Table S5. Transcription factors downstream of ERK activation likely to regulate genes in clusters 2,3, and 4.

Supplemental Table S6. MAGMA analysis of 28 known AD risk genes differentially expressed in NanoString dataset.

Supplemental Table S7. Human frontal cortex proteomics from AD, AD, AD/PD and control brains.

Supplemental Table S8. Differentially expressed ERK, JNK, and p38 phosphopeptides from human frontal cortex samples.

Supplemental Table S9. Upstream regulators of differentially expressed MAPK phosphopeptides in AD.

## REFERENCES

Arkun Y, Yasemi M (2018) Dynamics and control of the ERK signaling pathway: Sensitivity, bistability, and oscillations. PLoS One 13: e0195513

Bachiller S, Jimenez-Ferrer I, Paulus A, Yang Y, Swanberg M, Deierborg T, Boza-Serrano A (2018) Microglia in Neurological Diseases: A Road Map to Brain-Disease Dependent-Inflammatory Response. Front Cell Neurosci 12: 488

Block ML, Zecca L, Hong JS (2007) Microglia-mediated neurotoxicity: uncovering the molecular mechanisms. Nat Rev Neurosci 8: 57–69

Cargnello M, Roux PP (2011) Activation and function of the MAPKs and their substrates, the MAPK-activated protein kinases. Microbiol Mol Biol Rev 75: 50–83

Dai J, Johnson ECB, Dammer EB, Duong DM, Gearing M, Lah JJ, Levey AI, Wingo TS, Seyfried NT (2018) Effects of APOE Genotype on Brain Proteomic Network and Cell Type Changes in Alzheimer’s Disease. Front Mol Neurosci 11

de la Cova C, Townley R, Regot S, Greenwald I (2017) A Real-Time Biosensor for ERK Activity Reveals Signaling Dynamics during C. elegans Cell Fate Specification. Dev Cell 42: 542–553.e544

de Leeuw CA, Mooij JM, Heskes T, Posthuma D (2015) MAGMA: Generalized Gene-Set Analysis of GWAS Data. PLOS Computational Biology 11: e1004219

Dubbelaar ML, Kracht L, Eggen BJL, Boddeke E (2018) The Kaleidoscope of Microglial Phenotypes. Front Immunol 9: 1753

Fourgeaud L, Través PG, Tufail Y, Leal-Bailey H, Lew ED, Burrola PG, Callaway P, Zagórska A, Rothlin CV, Nimmerjahn A et al (2016) TAM receptors regulate multiple features of microglial physiology. Nature 532: 240–244

Gao T, Jernigan J, Raza SA, Dammer EB, Xiao H, Seyfried NT, Levey AI, Rangaraju S (2019) Transcriptional regulation of homeostatic and disease-associated-microglial genes by IRF1, LXRβ, and CEBPα. Glia 67: 1958–1975

Geraghty AC, Gibson EM, Ghanem RA, Greene JJ, Ocampo A, Goldstein AK, Ni L, Yang T, Marton RM, Pasca SP et al (2019) Loss of Adaptive Myelination Contributes to Methotrexate Chemotherapy-Related Cognitive Impairment. Neuron 103: 250–265 e258

Goetz EM, Ghandi M, Treacy DJ, Wagle N, Garraway LA (2014) ERK Mutations Confer Resistance to Mitogen-Activated Protein Kinase Pathway Inhibitors. Cancer Research 74: 7079–7089

Guo L, Bertola DR, Takanohashi A, Saito A, Segawa Y, Yokota T, Ishibashi S, Nishida Y, Yamamoto GL, Franco JFdS, et al (2019) Bi-allelic *CSF1R* Mutations Cause Skeletal Dysplasia of Dysosteosclerosis-Pyle Disease Spectrum and Degenerative Encephalopathy with Brain Malformation. The American Journal of Human Genetics 104: 925–935

Hagemeyer N, Hanft KM, Akriditou MA, Unger N, Park ES, Stanley ER, Staszewski O, Dimou L, Prinz M (2017) Microglia contribute to normal myelinogenesis and to oligodendrocyte progenitor maintenance during adulthood. Acta Neuropathol 134: 441–458

Harvey CD, Ehrhardt AG, Cellurale C, Zhong H, Yasuda R, Davis RJ, Svoboda K (2008) A genetically encoded fluorescent sensor of ERK activity. Proceedings of the National Academy of Sciences 105: 19264–19269

Hickman S, Izzy S, Sen P, Morsett L, El Khoury J (2018) Microglia in neurodegeneration. Nat Neurosci 21: 1359–1369

Hutton SR, Otis JM, Kim EM, Lamsal Y, Stuber GD, Snider WD (2017) ERK/MAPK Signaling Is Required for Pathway-Specific Striatal Motor Functions. The Journal of Neuroscience 37: 8102–8115

Keren-Shaul H, Spinrad A, Weiner A, Matcovitch-Natan O, Dvir-Szternfeld R, Ulland TK, David E, Baruch K, Lara-Astaiso D, Toth B et al (2017) A Unique Microglia Type Associated with Restricting Development of Alzheimer’s Disease. Cell 169: 1276–1290.e1217

Kim SH, Smith CJ, Van Eldik LJ (2004) Importance of MAPK pathways for microglial pro-inflammatory cytokine IL-1 beta production. Neurobiol Aging 25: 431–439

Konishi Y, Terai K, Furuta Y, Kiyonari H, Abe T, Ueda Y, Kinashi T, Hamazaki Y, Takaori-Kondo A, Matsuda M (2018) Live-Cell FRET Imaging Reveals a Role of Extracellular Signal-Regulated Kinase Activity Dynamics in Thymocyte Motility. iScience 10: 98–113

Konno T, Kasanuki K, Ikeuchi T, Dickson DW, Wszolek ZK (2018) *CSF1R*-related leukoencephalopathy. A major player in primary microgliopathies 91: 1092–1104

Krasemann S, Madore C, Cialic R, Baufeld C, Calcagno N, El Fatimy R, Beckers L, O’Loughlin E, Xu Y, Fanek Z et al (2017) The TREM2-APOE Pathway Drives the Transcriptional Phenotype of Dysfunctional Microglia in Neurodegenerative Diseases. Immunity 47: 566–581.e569

Lambert JC, Ibrahim-Verbaas CA, Harold D, Naj AC, Sims R, Bellenguez C, DeStafano AL, Bis JC, Beecham GW, Grenier-Boley B et al (2013) Meta-analysis of 74,046 individuals identifies 11 new susceptibility loci for Alzheimer’s disease. Nat Genet 45: 1452–1458

Liddelow SA, Guttenplan KA, Clarke LE, Bennett FC, Bohlen CJ, Schirmer L, Bennett ML, Munch AE, Chung WS, Peterson TC et al (2017) Neurotoxic reactive astrocytes are induced by activated microglia. Nature 541: 481–487

Morris EJ, Jha S, Restaino CR, Dayananth P, Zhu H, Cooper A, Carr D, Deng Y, Jin W, Black S et al (2013) Discovery of a Novel ERK Inhibitor with Activity in Models of Acquired Resistance to BRAF and MEK Inhibitors. Cancer Discovery 3: 742–750

Namsi A, Nury T, Hamdouni H, Yammine A, Vejux A, Vervandier-Fasseur D, Latruffe N, Masmoudi-Kouki O, Lizard G (2018) Induction of Neuronal Differentiation of Murine N2a Cells by Two Polyphenols Present in the Mediterranean Diet Mimicking Neurotrophins Activities: Resveratrol and Apigenin. Diseases 6: 67

Oosterhof N, Chang IJ, Karimiani EG, Kuil LE, Jensen DM, Daza R, Young E, Astle L, van der Linde HC, Shivaram GM et al (2019) Homozygous Mutations in CSF1R Cause a Pediatric-Onset Leukoencephalopathy and Can Result in Congenital Absence of Microglia. The American Journal of Human Genetics 104: 936–947

Ping L, Duong DM, Yin L, Gearing M, Lah JJ, Levey AI, Seyfried NT (2018) Global quantitative analysis of the human brain proteome in Alzheimer’s and Parkinson’s Disease. Sci Data 5: 180036

Rajendran L, Paolicelli RC (2018) Microglia-Mediated Synapse Loss in Alzheimer’s Disease. The Journal of Neuroscience 38: 2911–2919

Rangaraju S, Dammer EB, Raza SA, Gao T, Xiao H, Betarbet R, Duong DM, Webster JA, Hales CM, Lah JJ et al (2018a) Quantitative proteomics of acutely-isolated mouse microglia identifies novel immune Alzheimer’s disease-related proteins. Mol Neurodegener 13: 34

Rangaraju S, Dammer EB, Raza SA, Rathakrishnan P, Xiao H, Gao T, Duong DM, Pennington MW, Lah JJ, Seyfried NT et al (2018b) Identification and therapeutic modulation of a pro-inflammatory subset of disease-associated-microglia in Alzheimer’s disease. Mol Neurodegener 13: 24

Rustenhoven J, Smith AM, Smyth LC, Jansson D, Scotter EL, Swanson MEV, Aalderink M, Coppieters N, Narayan P, Handley R et al (2018) PU.1 regulates Alzheimer’s disease-associated genes in primary human microglia. Mol Neurodegener 13: 44–44

Santos SD, Verveer PJ, Bastiaens PI (2007) Growth factor-induced MAPK network topology shapes Erk response determining PC-12 cell fate. Nat Cell Biol 9: 324–330

Sellgren CM, Gracias J, Watmuff B, Biag JD, Thanos JM, Whittredge PB, Fu T, Worringer K, Brown HE, Wang J et al (2019) Increased synapse elimination by microglia in schizophrenia patient-derived models of synaptic pruning. Nat Neurosci 22: 374–385

Seyfried NT, Dammer EB, Swarup V, Nandakumar D, Duong DM, Yin L, Deng Q, Nguyen T, Hales CM, Wingo T et al (2017) A Multi-network Approach Identifies Protein-Specific Co-expression in Asymptomatic and Symptomatic Alzheimer’s Disease. Cell Syst 4: 60–72.e64

Shih R-H, Wang C-Y, Yang C-M (2015) NF-kappaB Signaling Pathways in Neurological Inflammation: A Mini Review. Front Mol Neurosci 8: 77–77

Silingardi D, Angelucci A, De Pasquale R, Borsotti M, Squitieri G, Brambilla R, Putignano E, Berardi N (2011) ERK Pathway Activation Bidirectionally Affects Visual Recognition Memory and Synaptic Plasticity in the Perirhinal Cortex. Frontiers in Behavioral Neuroscience 5

Smith AM, Gibbons HM, Oldfield RL, Bergin PM, Mee EW, Faull RLM, Dragunow M (2013) The transcription factor PU.1 is critical for viability and function of human brain microglia. Glia 61: 929–942

Spangenberg E, Severson PL, Hohsfield LA, Crapser J, Zhang J, Burton EA, Zhang Y, Spevak W, Lin J, Phan NY et al (2019) Sustained microglial depletion with CSF1R inhibitor impairs parenchymal plaque development in an Alzheimer’s disease model. Nature Communications 10: 3758

Tay TL, Mai D, Dautzenberg J, Fernandez-Klett F, Lin G, Sagar, Datta M, Drougard A, Stempfl T, Ardura-Fabregat A et al (2017) A new fate mapping system reveals context-dependent random or clonal expansion of microglia. Nat Neurosci 20: 793–803

Wood LB, Winslow AR, Proctor EA, McGuone D, Mordes DA, Frosch MP, Hyman BT, Lauffenburger DA, Haigis KM (2015) Identification of neurotoxic cytokines by profiling Alzheimer’s disease tissues and neuron culture viability screening. Sci Rep 5: 16622–16622

Yates B, Braschi B, Gray KA, Seal RL, Tweedie S, Bruford EA (2017) Genenames.org: the HGNC and VGNC resources in 2017. Nucleic Acids Res 45: D619–D625

Yin Z, Raj D, Saiepour N, Van Dam D, Brouwer N, Holtman IR, Eggen BJL, Möller T, Tamm JA, Abdourahman A et al (2017) Immune hyperreactivity of Aβ plaque-associated microglia in Alzheimer’s disease. Neurobiology of Aging 55: 115–122

Zambon AC, Gaj S, Ho I, Hanspers K, Vranizan K, Evelo CT, Conklin BR, Pico AR, Salomonis N (2012) GO-Elite: a flexible solution for pathway and ontology over-representation. Bioinformatics (Oxford, England) 28: 2209–2210

Zhang W, Liu HT (2002) MAPK signal pathways in the regulation of cell proliferation in mammalian cells. Cell Res 12: 9–18

Zhang Y, Chen K, Sloan SA, Bennett ML, Scholze AR, O’Keeffe S, Phatnani HP, Guarnieri P, Caneda C, Ruderisch N et al (2014) An RNA-sequencing transcriptome and splicing database of glia, neurons, and vascular cells of the cerebral cortex. J Neurosci 34: 11929–11947

Zheng C, Zhou X-W, Wang J-Z (2016) The dual roles of cytokines in Alzheimer’s disease: update on interleukins, TNF-α, TGF-β and IFN-γ. Translational Neurodegeneration 5: 7

