## Supplemental Figures for "Microglial ERK signaling is a critical regulator of pro-inflammatory immune responses in Alzheimer’s disease"

Supplemental figures: 4

Supplemental tables: 9

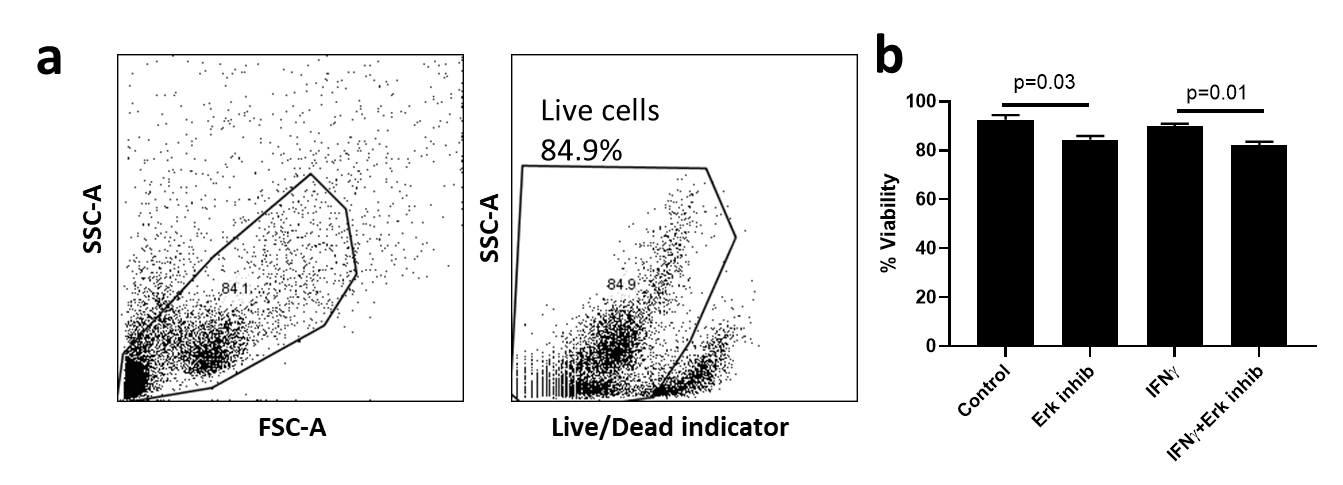

**Supplemental Figure S1. ERK inhibition exerts minimal impact on viability of microglia.**

(a) Flow cytometry gating strategy to identify live cell populations.

(b) Bar plot showing the slight reduction of viability among ERK inhibited microglia samples. N=3 replicates per group. Error bars represent SEM.

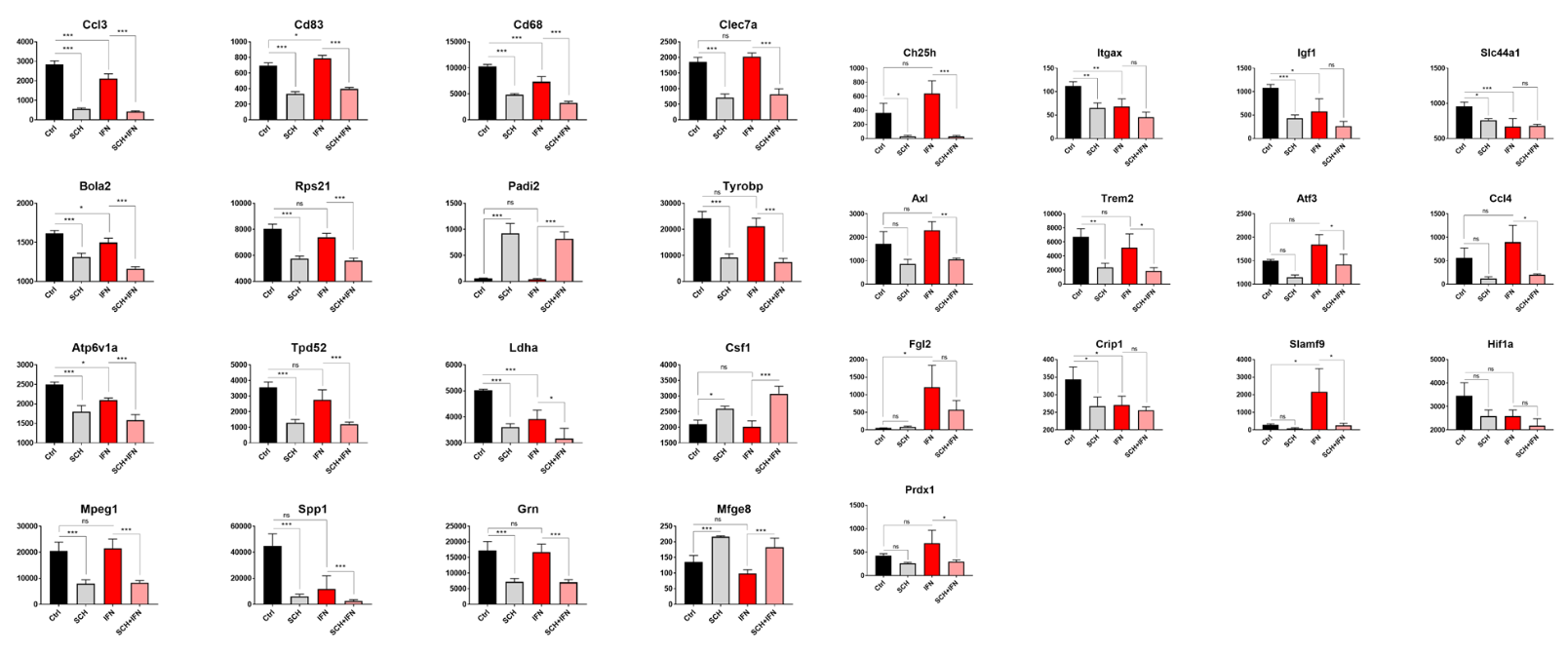

**Supplemental Figure S2. DAM genes positively regulated by ERK signaling.**

Individual NanoString normalized counts data for canonical DAM genes previously identified by single-cell RNA seq (Keren-Shaul et al, 2017) N=3 replicates per group. Error bars represent SEM. *p<0.05, **p<0.01, ***p<0.005.

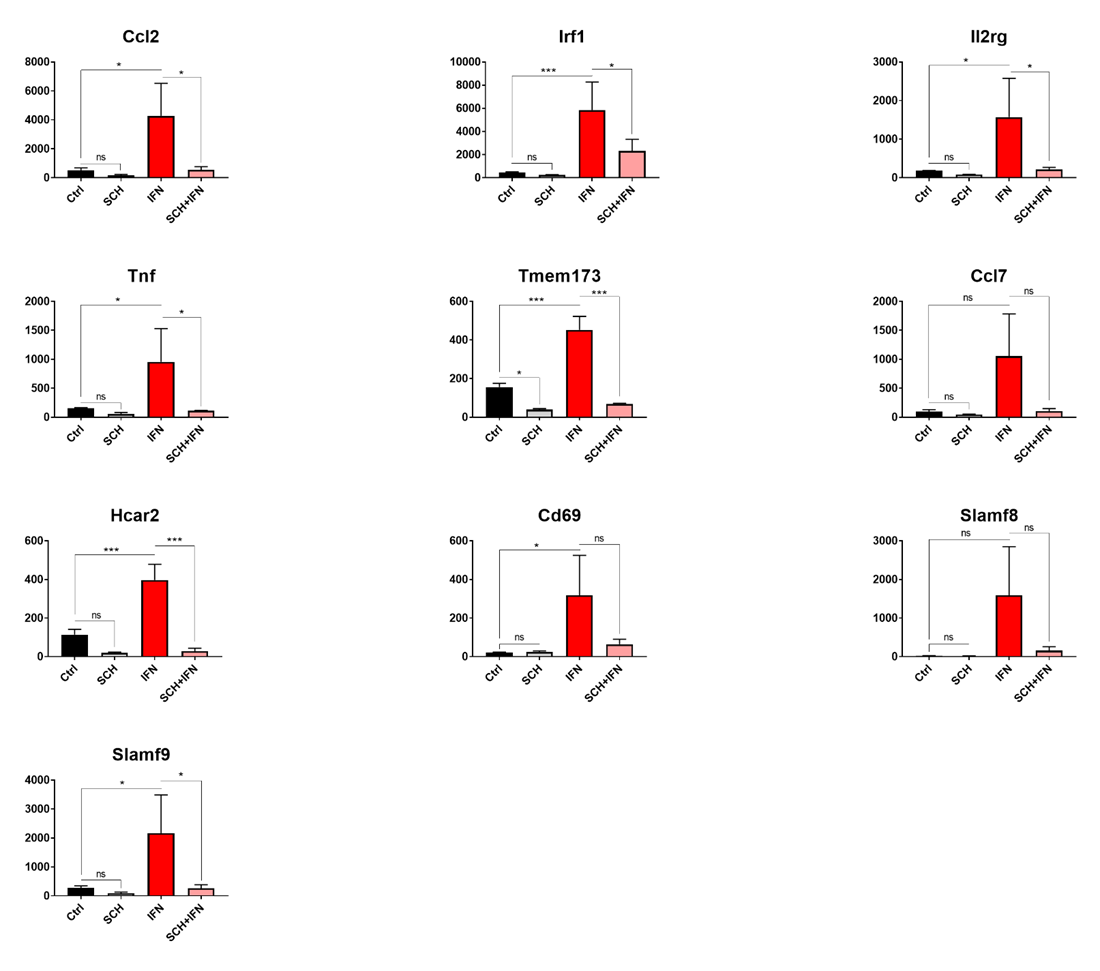

**Supplemental Figure S3. Pro-inflammatory DAM genes positively regulated by ERK signaling.** Individual NanoString normalized counts data for pro-inflammatory DAM genes previously identified by network analyses of microglial transcriptomes (Rangaraju et al, 2018b). N=3 replicates per group. Error bars represent SEM. *p<0.05, **p<0.01, ***p<0.005.

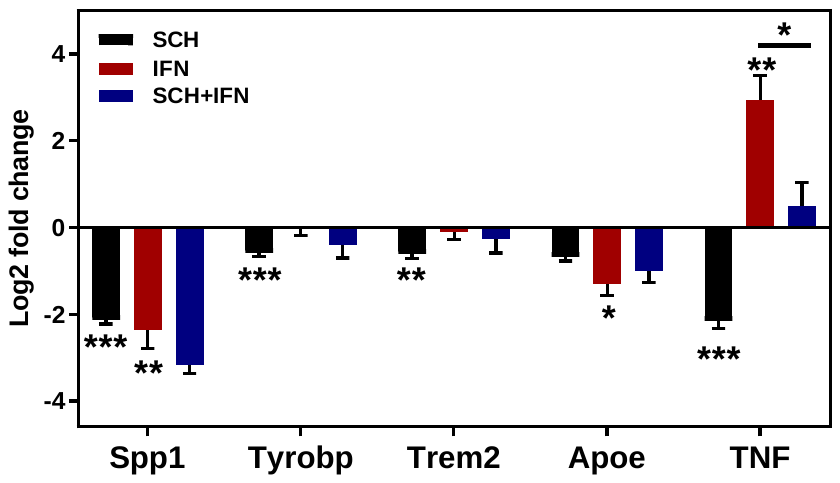

**Supplemental Figure S4. qRT-PCR validation confirms ERK effect of select DAM genes.**

Relative gene expression normalized to the housekeeping gene GAPDH (2-ΔΔCt method) is shown. N=3 replicates per group. Error bars represent SEM. *p<0.05, **p<0.01, ***p<0.005.
